# Oncogenic signaling in the adult *Drosophila* prostate-like accessory gland leads to activation of a conserved pro-tumorigenic program, in the absence of proliferation

**DOI:** 10.1101/2024.05.10.593549

**Authors:** S. Jaimian Church, Ajai J. Pulianmackal, Joseph A. Dixon, Luke V. Loftus, Sarah R. Amend, Kenneth Pienta, Frank C. Cackowski, Laura A. Buttitta

## Abstract

*Drosophila* models for tumorigenesis and metastasis have revealed conserved mechanisms of signaling that are also involved in mammalian cancer. Many of these models use the proliferating tissues of the larval stages of *Drosophila* development, when tissues are highly mitotically active, or stem cells are abundant. Fewer *Drosophila* tumorigenesis models use adult animals to initiate tumor formation when many tissues are largely terminally differentiated and postmitotic. The *Drosophila* accessory glands are prostate-like tissues and a model for some aspects of prostate tumorigenesis using this tissue has been explored. In this model, oncogenic signaling was induced during the proliferative stage of accessory gland development, raising the question of how oncogenic activity would impact the terminally differentiated and postmitotic adult tissue. Here, we show that oncogenic signaling in the adult *Drosophila* accessory gland leads to activation of a conserved pro-tumorigenic program, similar to that observed in mitotic larval tissues, but in the absence of proliferation. Oncogenic signaling in the adult postmitotic gland leads to tissue hyperplasia with nuclear anaplasia and aneuploidy through endoreduplication, which increases polyploidy and occasionally results in non-mitotic neoplastic-like extrusions. We compare gene expression changes in our *Drosophila* model with that of endocycling prostate cancer cells induced by chemotherapy, which potentially mediate tumor recurrence after treatment. Similar signaling pathways are activated in the *Drosophila* gland and endocycling cancer cells, suggesting the adult accessory glands provide a useful model for aspects of prostate cancer progression that do not involve cellular proliferation.

## Introduction

The fly *Drosophila melanogaster* has proven to be a convenient and inexpensive animal to model aspects of many human diseases, including tumorigenesis and cancer (Gong et al., 2021; Ugur et al., 2016). Their short lifespan, low cost of maintenance and the conservation of many tumorigenic signaling pathways, have led to a robust field of study using *Drosophila* models to investigate the cellular biology and genetics of tumorigenesis across tissues and developmental stages (Dow and Romero, 2010; Moraes and Montagne, 2021; Verheyen, 2022). This is facilitated by tools that allow spatio-temporal control over genetic manipulations including inducible over-expression or gene knockdown through RNAi as well as mosaic loss-of-function and gain-of-function tools (Bangi et al., 2016; Harnish et al., 2021; Sonoshita and Cagan, 2017; Zirin et al., 2020).

Prostate cancer is the second most diagnosed cancer for men worldwide and the third leading cause of cancer-related death for men over 65 (Culp et al., 2020). Half of men between 70-80 years of age show histological evidence of malignancy in the prostate (Carter et al., 1990). The *Drosophila* accessory glands serve reproductive functions analogous to the mammalian prostate, raising the possibility that these tissues in aged adult animals could be useful models for prostate cancer (Corrigan et al., 2014; Ito et al., 2014; Leiblich et al., 2019; Rambur et al., 2020; Rambur et al., 2021; Wilson et al., 2017). The fly reproductive system contains two accessory glands that release seminal fluid components into the Ejaculatory duct, where the seminal vesicles and testes also deposit sperm and additional seminal fluid components (Adams and Wolfner, 2007; Bloch Qazi and Wolfner, 2003; Cridland et al., 2022; Heifetz et al., 2005; Lung et al., 2001; Ravi Ram et al., 2005; Schnakenberg et al., 2012; Wilson et al., 2017). The fly accessory glands (AGs) have a similar, although simpler epithelial organization to the mammalian prostate. The accessory glands are composed of a polarized secretory epithelial monolayer that forms an apical lumen, surrounded on the basal side by extracellular matrix and a contractile muscle sheath (De Marzo et al., 2010; Rambur et al., 2021; Wilson et al., 2017). The secretory epithelial cells consist of two cell types termed main cells and secondary cells. Main cells comprise the bulk of the secretory epithelium making up about 1000 cells per gland lobe, while approximately 40-50 large highly secretory secondary cells are located at the distal tip of each gland (Bairati, 1968; Bertram et al., 1992; Sitnik et al., 2016; Susic-Jung et al., 2012; Takashima et al., 2023). The main and secondary cells in the mature adult accessory gland are believed to be largely quiescent. Both of these cell types are known to be polyploid and bi-nucleate due to variant, truncated cell cycles lacking mitosis that occur during pupal development (Bairati, 1968; Taniguchi et al., 2014; Taniguchi et al., 2018). Recent studies have revealed that there are additional adult-specific endoreplication cell cycles lacking mitoses very early post-eclosion and in secondary cells after mating (Box et al., 2024; Leiblich et al., 2019; Sekar et al., 2023). However, unlike the mammalian prostate, this tissue is thought to be completely postmitotic in adults with no known proliferating or stem cell population.

Prior studies in the AG have established that these tissues can be used to model important aspects of tumorigenesis. Cell growth and migration in the gland are impacted by genes that also play roles in human prostate cancer progression (Ito et al., 2014; Leiblich et al., 2019; Sekar et al., 2023). In addition, activation of Ras/Map Kinase and Insulin/PI3Kinase signaling, known pathways involved in prostate cancer, could lead to tissue hyperplasia with features of neoplastic basal tissue extrusions resembling early stages of metastases in this tissue (Rambur et al., 2020). However, in this study the genetic manipulations leading to pathway activation were performed during the pupal stages of development, specifically prior to the final mitoses and terminal differentiation in the developing accessory gland. Thus, it remained unclear which of the hyperplastic and neoplastic effects could be due to delays in cell cycle exit or defects in terminal differentiation prior to adulthood. This led us to wonder how the adult tissue would respond to oncogenic-like cell cycle activation after terminal differentiation and cell cycle exit.

Here, we investigated the effects of inducing cell cycle reactivation in the postmitotic adult accessory gland epithelium. Our goal was to investigate the level of cell cycle plasticity and capacity for cell cycle re-entry in accessory gland cells after they had entered quiescence and fully terminally differentiated. We find that adult accessory gland cells can re-enter a non-mitotic endoreduplication cell cycle, which leads to increased cellular hypertrophy and ploidy in the absence of mitosis. Extreme levels of cell cycle re-entry across many cells in the gland led to tissue hyperplasia and aneuploidy, activation of growth and stress signaling pathways, and apical and basal cellular extrusions, all in the absence of mitoses. Interestingly, transcriptomic analyses revealed a gene expression program that overlaps with other models of tumorigenesis in *Drosophila* in mitotically proliferating tissues, such as the larval imaginal tissues. This indicates that significant aspects of the aberrant signaling observed in these tumor models can occur in the absence of mitoses. We extended our analysis to investigate conserved pathways in non-dividing polyploid cells from mammalian prostate cancer models post-chemotherapy. Here, we observed that endocycling prostate cancer cells with increased genomic content (poly-aneuploid) exhibit similar gene expression changes in orthologous genes to those we observed in our *Drosophila* model of prostate-like hyperplasia. This suggests that *Drosophila* models of non-proliferating polyploid cells in the adult prostate-like tissue might contribute substantially to understanding prostate cancer cell states even in the absence of cellular proliferation.

## Results

### Exit from quiescence in the accessory gland induces epithelial hyperplasia via endocycles

The *Drosophila* accessory gland is composed of a single-layered, polarized secretory epithelium. The apical cell surface faces the gland lumen, while the basal surface is in contact with a collagen-containing basement membrane surrounded by a layer of muscle (Fig. 1A). The secretory epithelial cells of the gland are divided into two cell types: main and secondary cells (Fig. 1A). There are roughly 40 large secondary cells per gland, located at the distal tip (Fig. 1A). Main cells comprise the remaining ∼1000 secretory cells within each gland. Both cell types in the gland produce various proteins and other molecules that aid in fertility.

**Fig. 1.**
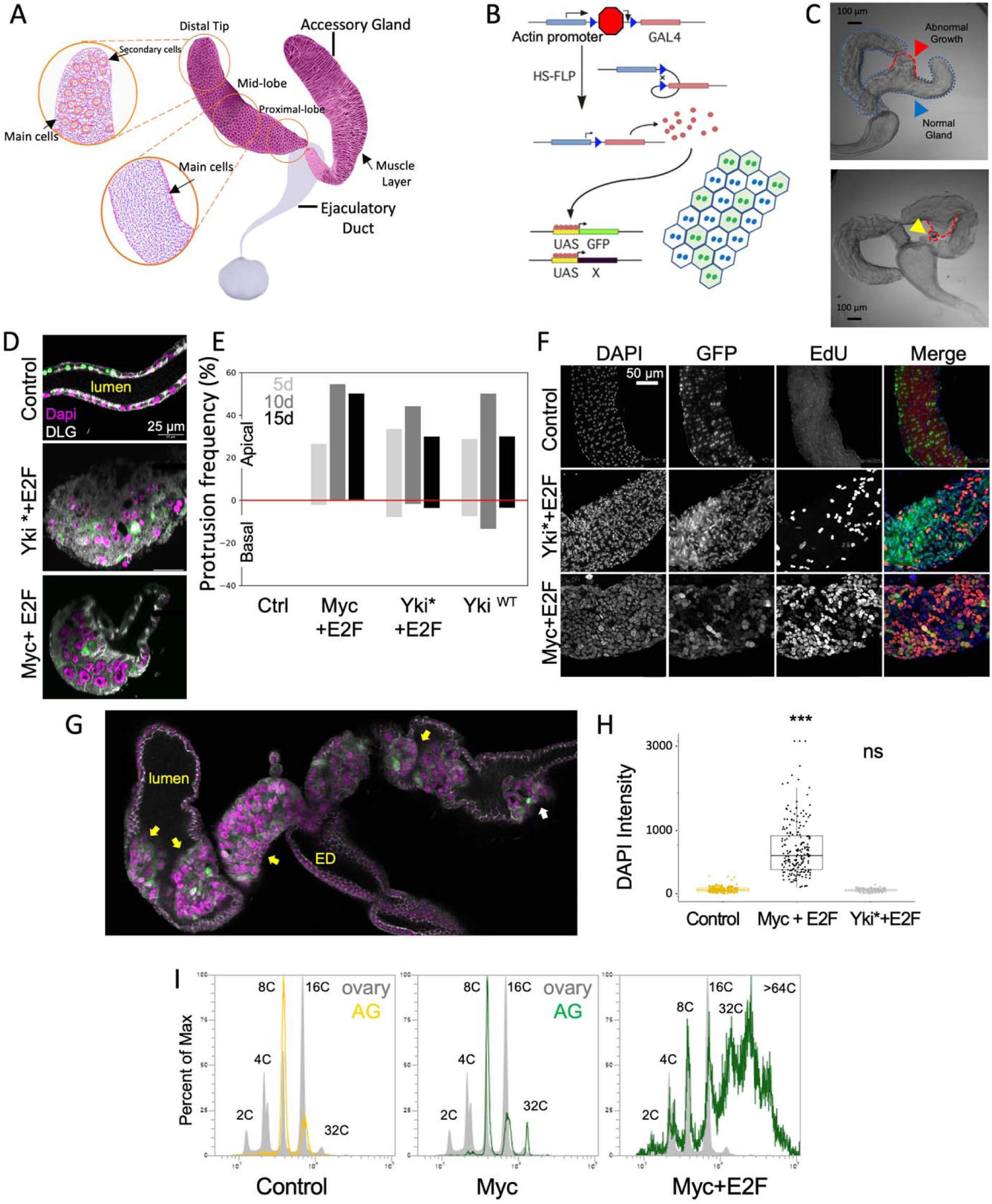
Oncogenic activation leads to endocycling and tissue hyperplasia in the adult accessory gland without proliferation. A) A cartoon diagram showing the major features of the *Drosophila* accessory gland. B) A diagram depicting the how transgenes are induced in our system via heat shock. C) DIC image of an accessory gland with protrusions when the Yki transgene is active. Abnormal basal hyperplasia and neoplasia like masses are present, outlined in red. D) Cross sections of accessory glands under control and oncogenic conditions. Oncogenic flies have apical growths invading the luminal space. E) Bar chart showing the frequency of accessory glands with apical or basal growths at various time points. Sample size: Control N= 254, Myc+E2f N= 276, Yki*+E2f N=161, and Yki-WT N=222. F) EdU assay of flies at ten days. All flies were heat-shocked for 20 minutes at 24-48h post eclosion and aged for six days before being fed 1mM EdU in 30% sucrose for 4 days before dissection. Sample size: Control N= 14, Myc+E2f N= 12, Yki*+E2f N=13. G) Dapi integrated density measurements (Dapi intensity) under oncogenic conditions. Sample size: 200 nuclei from 4 glands. H) Flow cytometry measurements of ploidy from isolated nuclei at 10 days. Grey are ploidy measurements from >20,000 ovary nuclei. At least 5,000 nuclei are represented in each histogram.

Although widespread genetic changes such as translocations and copy number variations are observed in the prostate, primary tumorigenesis is thought to arise clonally (Wang et al., 2018). As the adult *Drosophila* accessory gland epithelium is polyploid, bi-nucleate, postmitotic, and largely quiescent, we utilized a widely used heat-shock induced “flipout” system to induce transgene overexpression via the Gal4/UAS system in the AG (Fig. 1B). This approach uses a flippase-induced removal of a stop cassette between an actin promoter and Gal4 coding sequence, allowing UAS-driven gene expression to be activated postmitotically by a heat shock performed in a temporally controlled manner (Pignoni and Zipursky, 1997). The system is also sporadic and titratable, such that the number of cells expressing Gal4 increases with the length of the heat-shock. We tested several heat-shock parameters with a goal to induce consistent postmitotic expression in the adult accessory gland epithelia starting 24-48h post eclosion (Supp. Fig. 1). We found the number of accessory gland cells expressing Gal4 using this approach to be variable from animal to animal, and highly dependent upon the specific genetic background, with levels varying depending upon which UAS transgenes were expressed, even though all transgenes were crossed into the same hs-flipase background. To maximize consistency, all the work shown here used 20min. heat-shock condition, 24-48h post eclosion. We noted that binucleate main cells are often labeled with Gal4 in neighboring pairs of cells even though the induction event occurs post mitotically. This is because adult main cells retain ring canals to one neighbor, resulting in pairs of cells that share cytoplasm (Box et al., 2024). One limitation of this approach is the use of an actin promoter to drive Gal4 expression, which leads to the UAS-dependent expression of transgenes in other tissues throughout the adult.

By 4-6h post eclosion, the polyploid and bi-nucleate accessory gland epithelial cells exit the cell cycle with two octoploid nuclei (16C total cellular DNA content). Under normal conditions, we observe a low level of main cells (<1% per day) continuing to endocycle, generating a small population of cells with two 16C nuclei (32C total cellular DNA content), with most adult accessory gland epithelial cells in a long-term quiescent state by a few hours post-eclosion (Box et al., 2024). Additional endocycles can be driven in main cells by expression of positive cell cycle regulators such as CyclinD/Cdk4 or Cyclin E (Box et al., 2024; Molano-Fernandez et al., 2022), but in these instances, the Gal4 driver used expresses prior to eclosion. To test whether expression of positive cell cycle regulators could induce cell cycle re-entry in the quiescent adult gland, we overexpressed the *Drosophila* activator E2F complex, E2F1 and DP (hereafter referred to as E2F), a known driver of the endocycle (Zielke et al., 2011), or *Drosophila* Myc, a pro-growth regulator that promotes cell cycle entry and endocycling (Pierce et al., 2004). To prevent the elimination of aberrantly cycling cells, we also co-expressed the *Drosophila* apoptosis inhibitor, P35 (Hay et al., 1994). We observed minimal effects on overall gland size or morphology with E2F or Myc expression, although we did observe S-phases via (EdU) incorporation in a fraction of main cells (up to 5% of cells), demonstrating that quiescent main cells can be readily induced to re-enter the cell cycle in adults through ectopic Myc or E2F activity (Supp. Fig. 2). In an extreme example where most cells of the gland were overexpressing very high levels of Gal4 driving E2F, we did observe morphological evidence of gland overgrowth, suggesting that very high levels of cell cycle re-entry could drive tumor-like overgrowth in the *Drosophila* prostate-like tissue.

To drive more extreme cell cycle re-entry and overgrowth, we increased the expression of Yorkie (Yki^WT^) the fly ortholog of YAP/TAZ, the key transcriptional regulator downstream of the Hippo pathway responsible for controlling organ size (Dong et al., 2007). When Yki was overexpressed in the adult gland, we occasionally observed abnormal hyperplastic masses basally protruding from the gland (Fig. 1C). Additionally, hollow, cyst-like protrusions were occasionally observed, most often at the distal tips of basal protrusions (bottom panel, Fig. 1C). To see if this phenotype could be further increased by forcing additional cell cycle re-entry, we combined Myc and E2F expression simultaneously or combined overexpression of E2F with a constitutively active form of Yorkie, referred to as Yki* in our study (Oh and Irvine, 2009). When Myc+E2F or Yki*+E2F are expressed in the adult gland, we rarely observe hyperplastic masses basally protruding from the gland but very frequently observe apically extruding cell masses filling the gland lumen (Fig. 1D, G). We quantified the frequency of gland growth alterations under Myc+E2F, Yki*+E2F, and Yki^WT^ expressing conditions for 5-15 days (Fig. 1E). Regardless of genotype, most overgrowths invade the lumen apically, with 50% or more of animals across genotypes exhibiting apical invasion of hyperplastic cells. One limitation of our time course quantification is that we noted significant decreases in animal viability after 10 days in the genotypes overexpressing E2F. This suggests that the most affected animals may have died by day 10, possibly due to cell cycle disruptions across various tissues.

The additional tissue overgrowth apically and basally prompted us to examine glands for evidence of mitotic proliferation via staining for Phosphorylated Histone H3 Ser10 (PH3). In total, we examined over ∼150 samples across genotypes and timepoints and found no evidence of PH3 staining (Supp Fig. 3). By contrast, we observed extensive S-phases via a 4-day EdU labeling at day 10 in 70-100% of animals (Fig. 1F). At the ten-day time point, we also measured the DAPI nuclear intensity for 200 nuclei per sample from 4 different glands per genotype (Fig. 1H). As expected, we found evidence of increased DNA content in Myc+E2F conditions where EdU positive nuclei were visibly larger, while Yki*+E2F expressing nuclei appeared more normal (Fig. 1H). This demonstrates that Myc+E2F expression led to multiple rounds of endocycles during the assay, while Yki*+E2F expression induced widespread cell cycle re-entry but with more limited rounds of endocycling (Fig. 1H).

The distribution of Dapi intensities in Myc+E2F expressing nuclei suggested intermediate ploidies, consistent with aneuploidy. To confirm this, we performed flow cytometry on AG nuclei at 10-days post Gal4 induction (Fig. 1H). Ovaries were used as a ploidy “ladder” to place 2C, 4C, 8C, and 16C peaks using the stereotypical double peak at 4C (Bosco et al., 2007). In control 10d animals expressing only GFP and P35 we observed mostly octoploid nuclei with some 16C nuclei, consistent with recent work (Box et al., 2024), while in Myc+P35 conditions, we observed a small 32C population of nuclei, consistent with the consistent but limited EdU labeling observed in this condition (Fig. 1H, Supp. Fig. 2). By contrast, many nuclei from cells expressing Myc+E2F with P35 have a DNA content well beyond 32C and exhibit intermediate DNA content indicative of aneuploidy.

Altogether, our data demonstrates that quiescent bi-nucleate AG cells are capable of cell cycle re-entry into the endocycle, a cell cycle variant lacking mitosis, leading to overgrowth via tissue hyperplasia with apical or basal protrusions and occasional cyst-like extrusions.

### Hyperplasia in the AG affects epithelial organization

We noted that glands exhibiting overgrowth with apical and basal protrusions also exhibited altered localization of the septate junction protein, Discs Large (Dlg, Fig. 1D, Figure. 2A). We confirmed this alteration was not specific to Dlg by staining for a second septate junction protein Coracle (Cora), at 5, 10 and 15 days post Gal4 induction (Supp Fig. 4) (Fehon et al., 1994; Woods and Bryant, 1993). We observed significant mis-localization of both proteins after 10 days and confirmed mis-localization of Cora at 15 days using line scans of fluorescence intensity across cell-cell junctions. Cora levels at cell-cell junctions (peaks) were reduced, while non-junctional staining was increased (valleys) in Myc+E2F or Yki*+E2F expressing animals (Fig. 2B). We next examined whether the actin muscle layer was disrupted under hyperplastic conditions. Staining for actin using fluorophore conjugated phalloidin revealed increased gaps in the muscle fibers under Yki*+E2F or Myc+E2F conditions (Fig. 2A, yellow arrows). We observed one of the cyst-like basal protrusions poking through one of the gaps in the muscle layer, suggesting the cyst-like structures extrude basally outside of the epithelium (Fig. 2C). This is reminiscent of a neoplastic-like phenotype in the accessory gland previously reported under conditions of oncogenic Ras^V12^ expression during pupal developmental stages and into adulthood (Rambur et al., 2020). With oncogenic Ras over-expression, this phenotype was reported at a frequency of 20-80% of glands varying from experiment to experiment. In our conditions, we observe neoplastic-like extrusions much more rarely, in 2 - 4.5% of glands depending on timepoint and genotype (Fig. 2D), with the caveat mentioned previously that our most affected animals likely die at later time points due to actin-Gal4 driven expression in other tissues. This suggests that the developmental timing of the oncogenic induction may have an important impact on the penetrance of the neoplastic-like phenotype and that more mature, terminally differentiated accessory gland cells are more refractory to developing this phenotype, even under strong re-activation of the cell cycle. Consistent with this, when we activated Ras^V12^ expression in adult glands 24-48h post eclosion, we never observed any neoplastic basal extrusions (0/15 animals). This indicates that oncogenic Ras may not induce a neoplastic-like phenotype in the accessory gland unless the cells are in a state of mitotic proliferation prior to cell cycle exit.

**Fig. 2.**
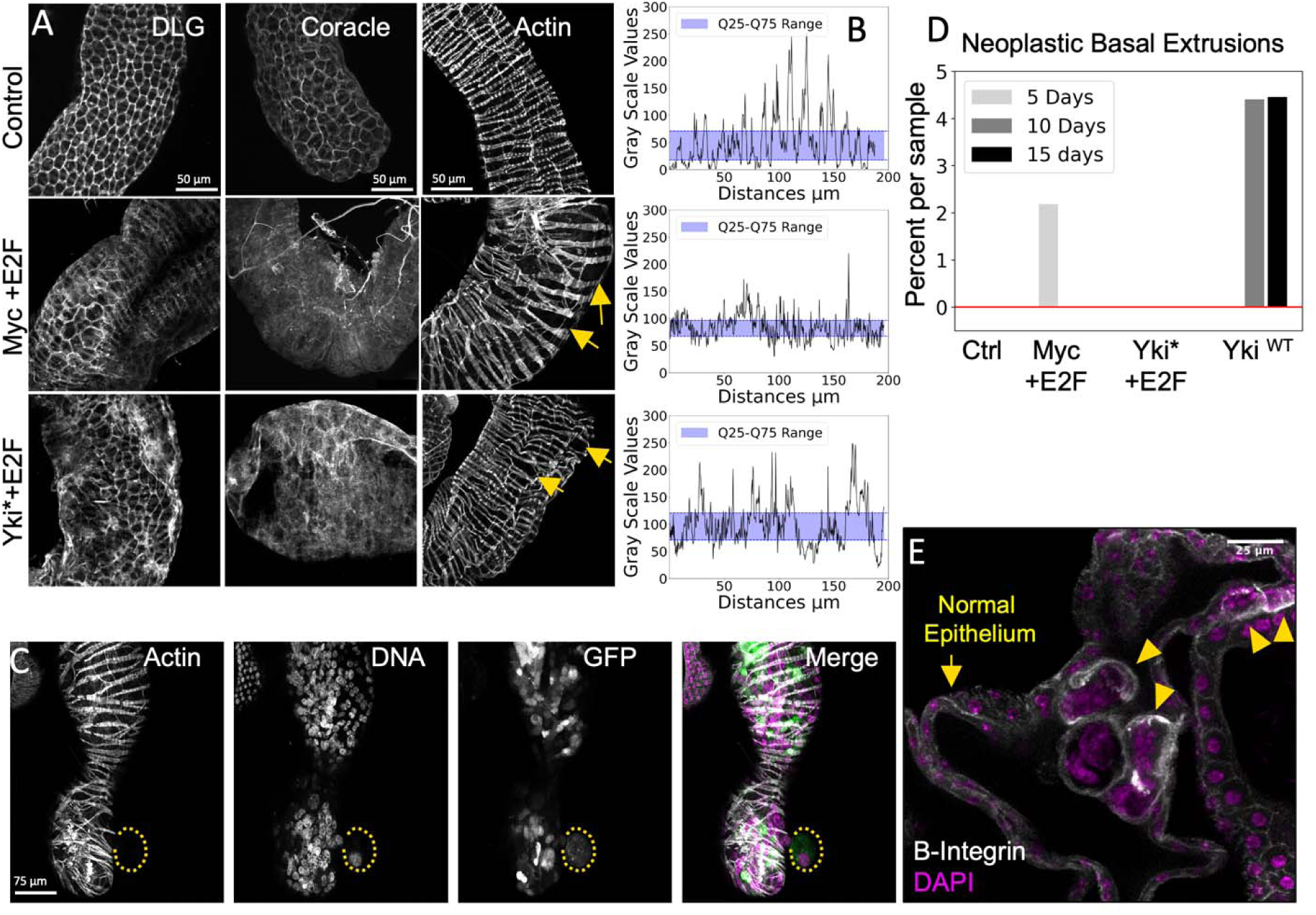
Hyperplasia in the adult accessory gland leads to disruption of cell adhesion and polarity with basal and apical extrusions. A) Septate junction protein localization and muscle organization under oncogenic conditions was assayed 15 days after heat shock. Septate junction proteins become mis-localized and gaps form in the muscle fibers surrounding the accessory gland (arrows). Sample sizes for DLG: Control N = 8, Myc+E2f N = 12, Yki*+E2f N = 6. Coracle: Control N = 6, Myc+E2f N = 11, Yki*+E2f N = 4. Actin/Phalloidin stain: Control N = 21, Myc+E2f N = 27, Yki*+E2f N = 15. B) Line scans of Coracle intensity for the images in panel A. The shaded blue area shows the interquartile range. C) A neoplastic-like mass in 5-day Myc+E2f expressing gland. The mass can be seen extruding basally in between muscle fibers. D) Bar chart of the frequency of neoplastic-like extrusions at different time points under the indicated genetic conditions. Sample size: Ctrl N = 311, Myc+E2F N = 299, Yki*+E2f N =180, Yki^WT^ N = 319. E) Z-section of the gland showing β integrin enrichment in 10-day old Myc+E2f expressing cells protruding apically into the lumen. Sample size: N =11

**Table 1.** Overlap of upregulated genes in oncogenic accessory glands and Drosophila tumor or wounding/regeneration literature.

| *Drosophila* tumor and wounding/regeneration literature | Log<sub>2</sub> Fold overlap* |
| --- | --- |
| The *Drosophila* Duox maturation factor is a key component of a positive feedback loop that sustains regeneration signaling | 2.67 |
| Promoter Proximal Pausing Limits Tumorous Growth Induced by the Yki Transcription Factor in *Drosophila* | 3.17 |
| An Ectopic Network of Transcription Factors Regulated by Hippo Signaling Drives Growth and Invasion of a Malignant Tumor Model | 2.25 |
| Interplay among *Drosophila* transcription factors Ets21c, Fos and Ftz-F1 drives JNK-mediated tumor malignancy | 2.83 |
| Shared enhancer gene regulatory networks between wound and oncogenic programs | 2.93 |
| Renal NF-κB activation impairs uric acid homeostasis to promote tumor-associated mortality independent of wasting | 3.74 |
| Cell polarity opposes Jak/STAT-mediated Escargot activation that drives intratumor heterogeneity in a *Drosophila* tumor model | 2.66 |

\*The log<sub>2</sub> fold of gene overlap, is shown for overlap 4-fold (log<sub>2</sub>2) or greater than that expected by chance (where chance is 1-fold, calculated using the hypergeometric distribution) for upregulated genes in our RNAseq dataset and the indicated published dataset.

### Neoplastic-Like vs. Hyperplastic-Like protrusions exhibit different signaling pathway activation

The presence of neoplastic extrusions and their association with basal protrusions led us to consider that apically or basally protruding cells may exhibit early metastatic phenotypes such as increased integrin expression and expression of matrix metalloproteinases to degrade the extracellular matrix. Indeed, we observed increased Beta-Integrin staining on some hyperplastic cells in both apical and basal extrusions (Fig. 2E and Supplemental Fig. 15) as well as increased MMP1 expression, which was present on many apical and basal protrusions but was particularly pronounced on neoplastic-like extrusions at the tip of basal protrusions (Fig. 3A and Supplemental Fig. 15). Myc+E2F clones exhibited a 3.6-fold increase in MMP1 staining compared to controls, and Yki*+E2F samples showed a 2.7-fold increase in MMP1 levels (Fig 3D).

**Fig. 3.**
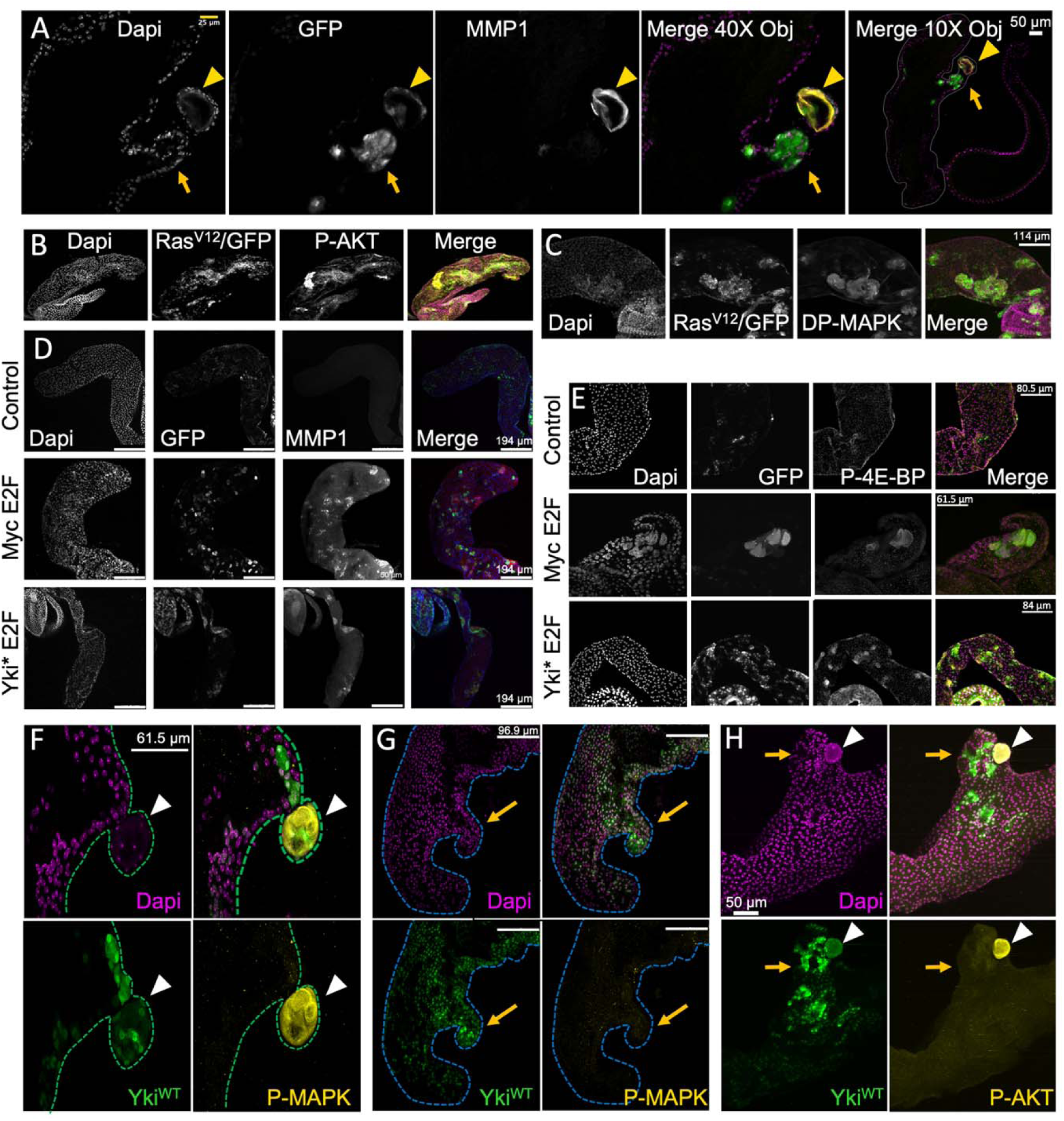
Neoplastic vs Hyperplastic growths exhibit different signaling pathway activity. A) A basal hyperplastic protrusion (arrow) with a neoplastic-like mass at the tip (arrowhead) positive for MMP1. B,C) Accessory glands overexpressing Ras^V12^ in GFP-labeled cells exhibit di-phosphorylated MAPK (DP-MapK) and phosphorylated AKT (pAKT) in the absence of hyperplasia or neoplasia at ten days. Sample size: N = 6 DP-MapK, N=2 pAKT. D) Oncogenic expression leads to increased MMP1 in 20d samples. Sample size: Control N=12, Myc+E2f, DP N=12, Yki*+E2f, Dp N= 13. E) Oncogenic expression leads to increased phospho-4EBP (p4EBP) in 10 d samples. Sample sizes: Control N=12, Myc+E2f, DP N=8, Yki*+E2f, Dp N= 10. F,G,H) Neoplastic-like masses (arrowheads) are positive for DP-MAPK (F) and P-AKT, while hyperplastic growths (arrows) are not (G,H) at 20d. Sample size: N= 14.

In our Yki^WT^ overexpression model, we observed the development of finger-like protrusions extending from the mid-lobe region of the accessory glands, as illustrated in (Fig. 3A). These structural anomalies resemble extra-prostatic extensions (EPE) observed in human prostate cancer associated with YAP1 overexpression (Collak et al., 2017). Notably, the tips of these protrusions were characterized by neoplastic-like masses (Fig. 3A). Further analysis involved MMP1 (Matrix Metallopeptidase 1) staining of an EPE to assess its expression levels. The results revealed that MMP1 staining in the neoplastic areas of the basal protrusions was significantly increased, showing a seven-fold enrichment compared to areas of hyperplasia (Fig. 3A).

We next compared the signaling pathways activated in our oncogenic contexts to those in cells expressing oncogenic hyperactive Ras^V12^ in neoplastic-like extrusions (Rambur et al., 2020). Despite effective activation of Map-Kinase signaling by Ras^V12^ overexpression, assayed through high levels of di-phosphorylated ERK (dp-ERK) and PI3K/AKT signaling, assayed by phosphorylated Akt (pAkt) antibody staining in adult tissues (Fig. B,C), we observed no basal protrusions or extrusions, and only moderate levels of cellular and nuclear hyperplasia when Ras^V12^ is activated in the adult gland after cell cycle exit. Consistent with our hypothesis that Ras^V12^ induced phenotypes differ after the gland is terminally differentiated, mature and quiescent.

We next examined whether the apical or basal protrusions or neoplastic-like growths induced by E2F+Myc, E2F+Yki* or Yki^WT^ expression exhibited PI3K/Akt and/or MAPK signaling activities similar to the neoplastic-like Ras-induced extrusions described previously. In all cases, apical or basal protrusions that did not have neoplastic extrusions were negative for high levels of MAPK signaling evident from dp-ERK, and PI3K induced phosphorylation of Akt (Supp Fig. 4). However, when these hyperplasias did exhibit neoplastic-like extrusions, the extrusions were positive for dp-ERK and pAkt (Fig. 3). Interestingly, all hyperplasias induced by E2F+Myc or E2F+Yki* were positive for phosphorylated-4EBP, a target of the TORC1 complex downstream of PI3K signaling, with or without extrusions, even though they were negative for pAkt (Fig. 3E, Supp Fig. 4). This suggests that hyperplastic growth in adult glands may engage overgrowth through PI3K/TORC1 signaling without activating the TORC2/Akt signaling branch (Frappaolo and Giansanti, 2023), unless additional events such as developmental proliferation or neoplastic extrusion with MAPK signaling also occur.

**Fig. 4.**
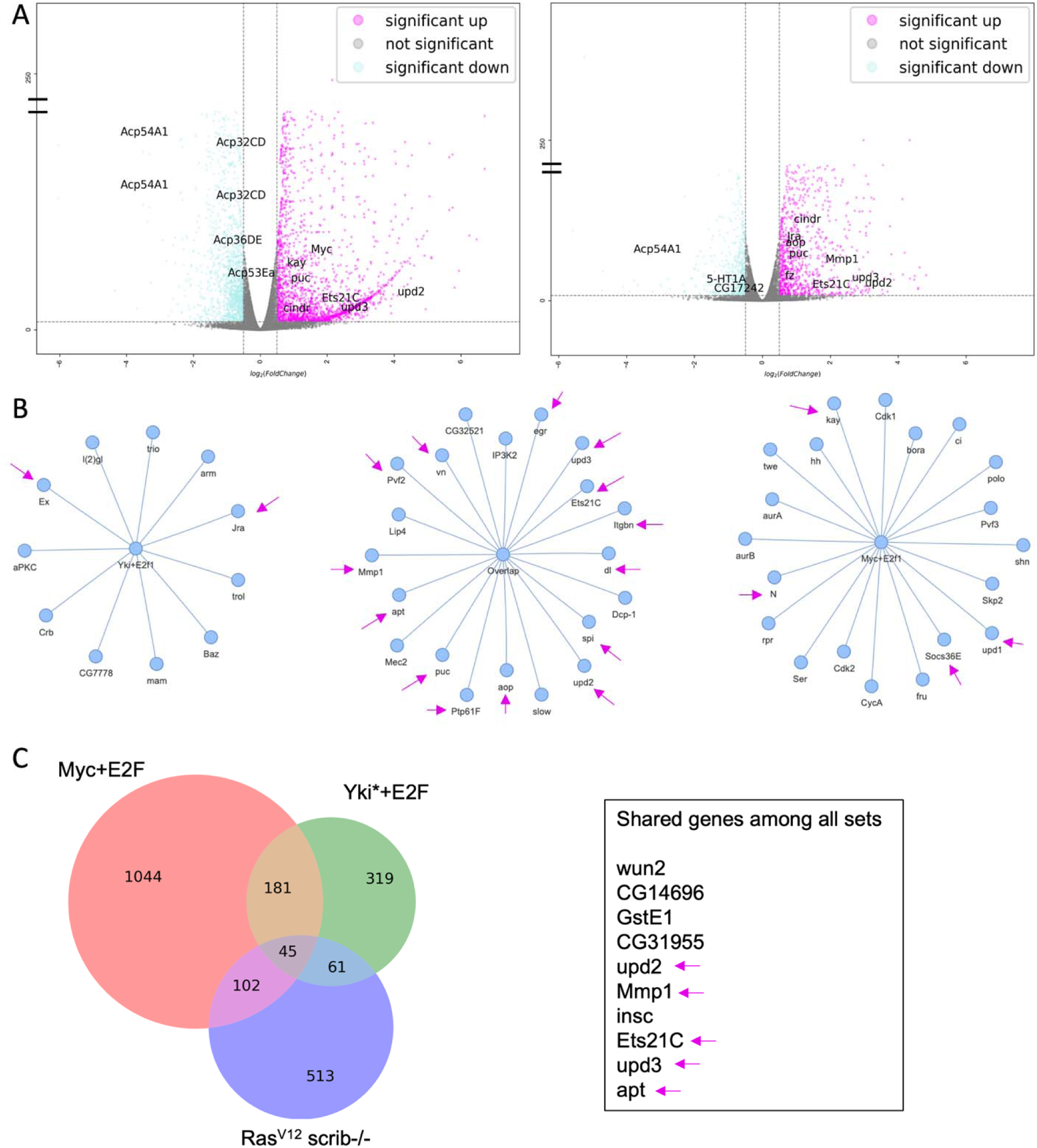
RNAseq reveals activation of a common tumorigenic program in the absence of mitoses. A) Volcano plots for Myc+E2f, Dp and Yki*+E2F, Dp show transcripts significantly up and downregulated. In both conditions genes associated with JNK signaling and proliferation networks were significantly upregulated while genes involved in male fertility were downregulated. B) Node networks of genes upregulated in *Drosophila* tumor models curated from the literature in Table 1. C) Overlap of genes upregulated in accessory glands expressing the indicated oncogenes with a commonly used *Ras,scrib* larval mitotic tumor model (Kulshammer et al., 2015). Shared genes with known roles in tumorigenic proliferation are indicated (arrows).

Further supporting the differences between hyperplastic growth and neoplastic-like extrusions, we found that neoplastic-like growths induced by Yki^WT^ in adult glands also exhibit high levels of dp-ERK and p-Akt, while Yki^WT^ -induced hyperplastic growths either with or without extruded neoplasias, did not exhibit high dp-ERK or p-Akt signal (Fig. 3 F-H). Importantly, 73% of the neoplasia-like masses we observe from adult activation of oncogenes developed from hyperplasia-like masses, while only 27% arose directly from non-hyperplastic epithelial tissue (compare Fig. 3 F,H to G). This suggests that neoplastic-like growths may initially be triggered by the same transcriptional signals that lead to hyperplasia in adult glands.

Based on this evidence, we suggest that the propensity for neoplastic-like behavior induced by oncogenic Ras depends on the initial cycling status of the prostate-like cells. If the cells are still proliferating, oncogenic Ras signaling can co-activate PI3K/Akt and Map Kinase signaling in a manner that results in a high frequency of neoplastic extrusions. By contrast, when the cells are quiescent in the adult, the activation of these signaling pathways by oncogenic Ras is no longer sufficient to lead to frequent neoplasia. Instead, cell cycle re-entry in adult cells can lead to hyperplastic growth that engages PI3K/TORC1 signaling, but not PI3K/Akt and MAPK signaling. More rarely in these conditions, additional events can lead to neoplastic conditions that aberrantly activate MAPK and PI3K/Akt signaling even in the absence of mitosis.

### A shared tumor signaling network is activated in hyperplastic tissues, even in the absence of proliferation

The expression of MMP1, Beta-integrin and phosphorylation of TOR targets suggested our Yki*+E2F, Yki^WT^, and Myc+E2F induced hyperplasias may be activating a shared tumorigenic program in the absence of mitotic proliferation. To test this, we performed RNA-Seq on our Yki*+E2F, Yki^WT^, and Myc+E2F, vs. control animals expressing GFP+P35 at the 10-day time point to identify shared gene expression changes. We visually confirmed similar levels of transgene expression using the UAS-GFP levels prior to RNA isolation for each sample (i.e. we selected glands with >⅓ of the cells GFP+). Each sample was comprised of 80-100 dissected glands with the ejaculatory duct removed, and we performed three independent replicates on polyA selected RNA sequencing. We observed a suite of upregulated and downregulated genes in each condition compared to the controls for immune response, cell cycle, responses to stress, and reproduction responses summarized in (Fig. 4A and Supplemental Fig. 6-9).

In both Yki*+E2F and Myc+E2F conditions, we observed upregulation of several genes known to be upregulated in wound healing and tumorigenic models in flies, including JNK signaling targets *Ets21C*, *MMP1*, *puckered* and hyperplasia-inducing Jak/STAT signaling pathway ligands *upd2* and *upd3* (Fig. 4A). We also observed the downregulation of accessory gland-specific genes (such as many Accessory Gland Proteins, Acps) and genes involved in fertility, mating behavior, and sperm competition (Fig. 4A). This suggests that hyperplasia and cell cycle re-entry occur at the expense of accessory gland function, (see downregulation of reproduction-associated genes in Fig. 4A and Supplemental Fig 9). The complete list of Gene Ontology (GO) terms enriched in up and downregulated gene sets is presented in (Supplemental Figures 10-11).

We observed a strong overlap between the Yki*+E2F and Myc+E2F gene expression signatures that converge on several genes known to be commonly mis-regulated in proliferative tumor models in various larval tissues, including imaginal discs and brains. We created a node-based graph depicting the common *Drosophila* tumor mis-regulated genes for in our oncogenic backgrounds, and the overlapping gene set between both datasets, (Fig. 4B). We also observe a stronger mis-regulation of cell cycle-related E2F targets (e.g. *cycA*, *cdk2*, *aurA*, *aurB, polo*) in the Myc+E2F signature that is not as strong in the Yki*+E2F gene expression signature (Fig. 4B). This is consistent with the positive feedback regulation between E2F and Myc and their known synergistic roles in regulating cell cycle entry from G1 (Duman-Scheel et al., 2004; Fernandez et al., 2003; Leone et al., 1997; Matsumura et al., 2003). By contrast, in the Yki*+E2F gene expression signature, we observe a stronger mis-regulation of genes involved in cell polarity, cell-cell adhesion complexes, and cytoskeletal regulators (e.g *arm*, *aPKC*, *Baz*, *Crb*, *Ex*) (Fig. 4B). This is also consistent with the known roles of the Hippo signaling pathway in regulating and responding to epithelial polarity, cellular adhesion, and tissue tension (Boggiano and Fehon, 2012; Borreguero-Munoz et al., 2019; Doggett et al., 2011; Elbediwy et al., 2016; Kroeger et al., 2024; Ling et al., 2010; Richardson and Portela, 2017; Sarpal et al., 2019; Su et al., 2017; Yang et al., 2015).

Our data suggests that hyperplasia without proliferation in the AG engages common tumorigenic gene networks described in other tumor models, even in the absence of mitosis (Hamaratoglu and Atkins, 2020; Khan et al., 2013; Logeay et al., 2022). In particular, we observed significant overlap with a common signature from the *Drosophila* metastatic tumor model of oncogenic Ras combined with the loss of epithelial polarity through loss of the Scribble protein (Ras^V12^,*scrib*-/- mutant) (Fig. 4C), (Atkins et al., 2016; Kulshammer et al., 2015) and wound healing gene expression profiles (Khan et al., 2017), Table 1). The overlap with wound healing and tumorigenic signatures encompasses genes involved in promoting proliferation (*Jra/Kay, Pvf, Trbl, upd*) as well as genes more recently described to be part of an aberrant “pro-senescence” program induced by JNK signaling present in wounding and tumorigenic models (*MMP1, Ptp61F, Socs36e,* (Floc’hlay et al., 2023). This demonstrates that genes involved in cancer hallmarks, such as cell invasion, loss of polarity, cellular hyperplasia, de-differentiation, and de-regulated protein synthesis, can all be simultaneously activated in the adult secretory accessory gland epithelium in the absence of proliferation. This suggests proliferation in the prostate-like epithelium is not required for these hallmarks.

### Overexpression of the ERG ortholog ETS21C drives cell cycle re-entry in the adult AG and synergizes with Myc to increase DNA replication

In our RNAseq study, the ERG ortholog and JNK target Ets21C was significantly upregulated in both of our Myc+E2F and Yki*+E2F hyperplasia models. Ets21C has previously been shown to transcriptionally activate cell cycle genes, including many E2F-regulated targets (Zhang et al., 2022). We therefore wondered whether Ets21C overexpression could drive cell cycle re-entry in the quiescent adult AG and how it compared to E2F or Myc overexpression.

A 4-day EdU labeling assay was conducted to assess transgene expression from days 6-10 across various conditions: Yki* without E2F, Myc without E2F, Ets21C, Ets21C+Myc, E2F, Myc+E2F, and Yki*+E2F (Fig. 5A). Observations indicated no or few EdU-labeled cells under control conditions (*hsflp* with only *UAS GFP* and *P35*) and for Yki* (Fig. 5B). Myc overexpression resulted in a modest yet statistically significant increase in EdU-labeled cells compared to the control, reflecting an increase from approximately 1 cell per 10 animals to an average of 1 cell per animal (Fig. 5B). By contrast, dramatically increased EdU labeling was observed in conditions expressing Ets21C, Myc+Ets21C, E2F, Myc+E2F, and Yki*+E2F. Notably, Myc+E2F and Yki*+E2F conditions achieved maximal labeling, with over 100 EdU+ cells per animal. When combined with Myc, both E2F and Ets21C exhibited an increased number of EdU cells per animal, with Myc+E2F showing statistical significance compared to the individual transgenes, p-value 0.002 (Fig. 5B), although Myc+Ets21C was not significantly different from Ets21C alone in the frequency of EdU+ cells.

**Fig. 5.**
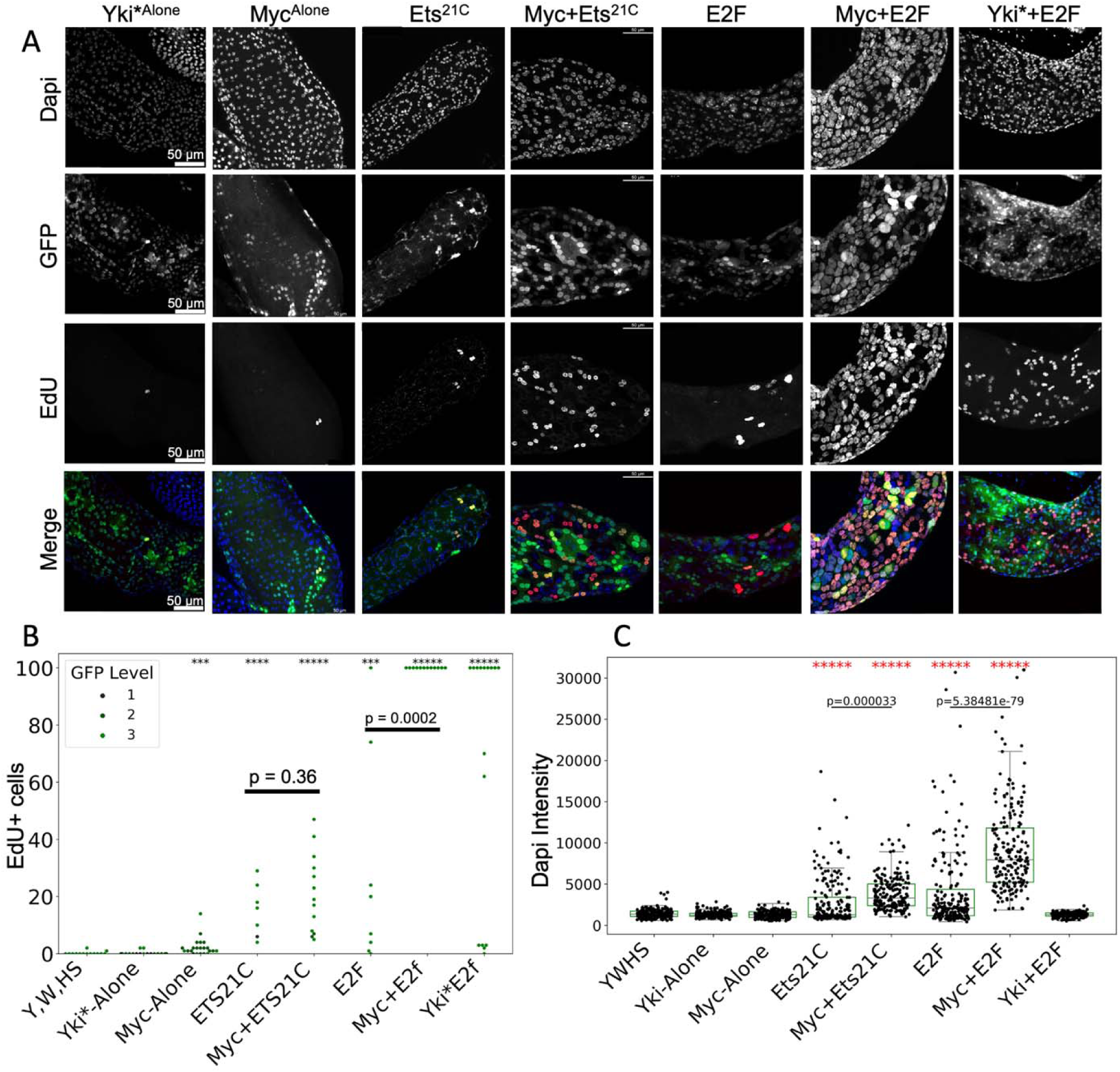
ETS21C overexpression in the accessory gland leads to increased DNA replication and aneuploidy. A) EdU assays show DNA replication from day 6-10 of expression of the indicated transgenes. Sample sizes: Control N= 14, Myc N = 22, Ets21C N =7, Myc+Ets21C N= 5, E2F N = 7, Myc+E2f N= 12, Yki*+E2f N=13, B) Quantification of EdU positive cells for the indicated genotypes, the dot color indicates the level of GFP expression with 3 being the strongest GFP. Samples with >100 EdU+ cells were plotted at a max of 100. Note nearly every sample from the E2F+Myc and E2F+Yki conditions has >100 labeled cells. C) Dapi intensity quantifications from 200 nuclei for 4 biological replicates for 10d samples of the indicated genotypes.

To further investigate the impact of Ets21C and E2F co-expression with Myc on ploidy, Dapi intensity was quantified from 200 nuclei across 4 biological replicates under the conditions described in Fig. 5A. Increased Dapi intensity was observed in Myc+E2F and Myc+Ets21C samples relative to controls as well as single transgene conditions (Fig. 5C). Myc+E2F and Myc+Ets21C models exhibited a continuous distribution of Dapi intensities, indicative of intermediate ploidies, suggestive of aneuploidy (Fig. 5C). This observation aligns with our previous findings for Myc+E2F. We suggest that as described in the *Drosophila* intestine, high levels of Ets21C expression act in a manner similar to high E2F activity to induce cell cycle genes involved in endocycling and DNA replication (Zhang et al., 2022), which is enhanced and potentiated with co-expression of Myc.

### High Ets21C promotes non-autonomous nuclear hyperplasia

We noted frequent non-autonomous effects on nearby cells in our experiments involving Ets21C overexpression as well as combinations of Myc+Ets21c and E2F or E2F+Myc, suggesting the hypertrophy and endoreplication driven by oncogenic signaling also impacted endocycling in nearby non-GFP expressing cells (Fig 6 A,C). To quantify this, we examined nuclear Dapi intensity for 100 GFP-positive cells and nearby GFP-negative cells across four biological replicates for overexpression of Ets21C, Myc+Ets21C, E2F, Myc+E2F. We observed increased nuclear areas across all genotypes for GFP+ cells with Ets21C expression showing a bi-modal distribution of nuclear areas and a long tail of some samples with very high integrated Dapi intensity, similar to E2F overexpression. When combined with Myc co-expression, Dapi intensity for GFP+ cells significantly and consistently increased, in agreement with the increased endocycling and EdU labeling we describe in Fig. 5. When we examine the distribution of nuclear Dapi intensities for GFP-negative cells neighboring GFP+ cells, we observe similarly increased nuclear size distributions, suggesting non-autonomous endocycling also occurs, which we also observed in our EdU labeling in Fig.5. Myc+Ets21C showed a statistically significant difference between the nuclear size of GFP+ and GFP-negative neighbors, with the non-autonomous endocycling being less than that of cells directly overexpressing Myc+Ets21C. With Myc+E2F co-expression, we observe dramatic endocycling along with dramatic endocycling of GFP-negative neighbor nuclei (Fig 6B,C) suggesting non-autonomous growth effects.

**Fig. 6.**
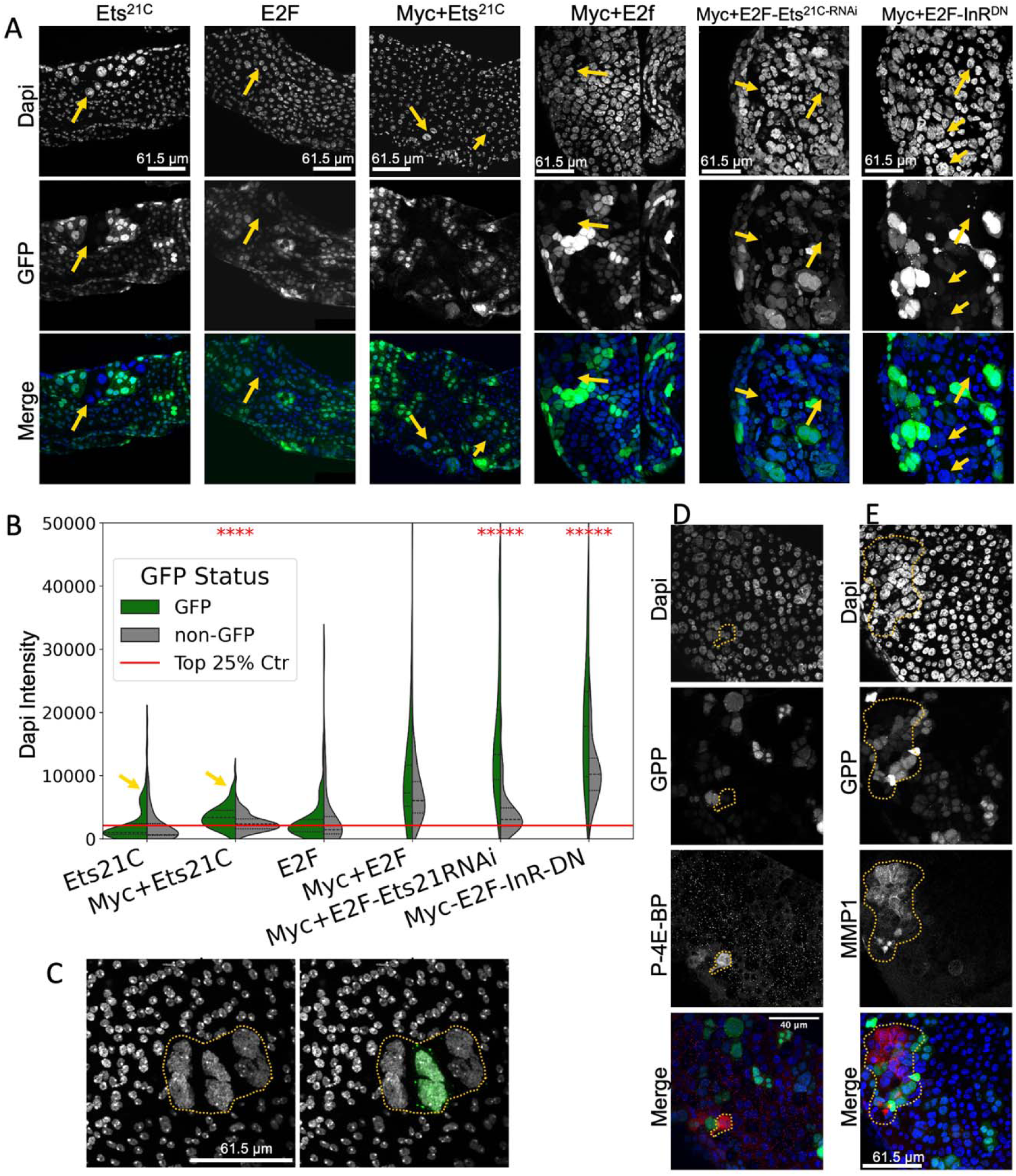
Ets21C impacts non-autonomous endocycling in GFP-negative neighboring cells. A) Examples of glands expressing Ets21C, E2F, Myc+Ets21C, Myc+E2F, Myc+Ets21C^RNAi^ and Myc+InR^DN^ in GFP-positive cells with Dapi in blue. Many GFP-negative cells exhibit nuclear hyperplasia indicative of non-autonomous induction of endocycling and cell growth (arrows). B) A split-violin plot of Dapi intensities for the indicated genotypes shown in panel A showing GFP-positive (green) vs. GFP-negative (grey) nearby cells. Yellow arrows indicate a subset of nuclei in Ets21C and Myc+Ets21C with very high DAPI intensity, like those shown in A. For each genotype, 100 GFP-positive and GFP-negative nuclei were quantified across 4 biological replicates. The red line indicates the upper quartile for the control (only P35 and GFP expression) Dapi intensity. C) An example of an Myc+E2F expressing sample with GFP-positive and GFP-negative nuclei exhibiting evidence of nuclear anaplasia. D,E) Examples of MMP1 (D) and p4EBP (E) staining in GFP-positive and GFP-negative cells of Myc+E2F expressing glands.

Non-autonomous effects of JNK signaling can be propagated through production of paracrine signaling factors, such as Upd ligands or diffusible reactive oxygen species, both of which can be triggered by aneuploidy (Clemente-Ruiz et al., 2016; Khan et al., 2017; La Fortezza et al., 2016; Muzzopappa et al., 2017; Santabarbara-Ruiz et al., 2015). Consistent with this, we observe non-autonomous activation of MMP1 expression in adjacent GFP-negative nuclei when Myc+E2F is used to induce tissue hyperplasia (Fig 6D). Ets21C which is induced by JNK has also been shown to activate expression of ligands that have non autonomous effects on growth such as *upd3, upd1, MMP1* and *Pvf1* (Mundorf et al., 2019; Toggweiler et al., 2016). We observed Myc+E2F-induced non-autonomous growth signaling and activation of the PI3Kinase pathway in adjacent GFP-negative nuclei, assessed via phosphorylation of 4E-BP, which could be mediated via Ets21C induction of targets or PI3K signaling (Fig 6E). We therefore next examined the effect of knocking down Ets21C using RNAi or co-expression of a dominant negative form of the insulin Receptor (InR^DN^) in the presence of Myc+E2F co-expression to determine roles for Ets21C and PI3K signaling in nuclear and cellular hyperplasia and non-autonomous nuclear and cellular hyperplasia. Knocking down Ets21C by RNAi with Myc+E2F co-expression had little effect on nuclear size in GFP+ cells but had a strong effect on reducing nuclear Dapi intensity in neighboring GFP-cells. By contrast co-expression of InR^DN^ in the presence of Myc+E2F did not significantly reduce nuclear sizes for GFP+ or GFP-negative neighboring cells, suggesting PI3K signaling may not be required to engage or sustain feedback loops that promote endocycling in this tissue. Alternatively, limiting InR signaling in other tissues may allow for increased endocycling induced by E2F+Myc in the accessory gland through tissue non-autonomous effects, or effects that improve animal survival. As our overexpression system results in UAS-target gene expression throughout the animal, we cannot rule out this possibility.

### Endocycling prostate cancer cells upregulate genes orthologous to those upregulated in hyperplastic *Drosophila* accessory glands

After chemotherapy, a minority of cancer cells in the treated population exit mitosis and undergo multiple rounds of endoreduplication. These cancer cells exhibit increased survival, hyperplasia, and increased genomic content, ie., they are in a large, polyaneuploid state (Kim et al., 2023; Lin et al., 2019; Pienta et al., 2021). These endocycling cells can also enter senescence-like states that can impact the tumor microenvironment in a non-autonomous manner (Lin et al., 2019; Niu et al., 2021). To examine whether our adult *Drosophila* accessory gland hyperplasia resembles the endocycling state in human prostate cancer cells, we induced polyploidy in a PC3 human prostate cancer cell line using docetaxel treatment. By three days of culture with 5nM Docetaxel, most PC3 cells die or arrest their cell cycle. Following our previously established protocols for endocycling cell formation (Kim et al., 2023; Mallin et al., 2023), we removed the docetaxel on day three, provided cells with fresh culture medium and tracked continued cellular growth for up to 45d in culture (Fig. 7A, B). We confirmed that at least a portion of this growth is due to endocycling, evident from EdU incorporation in S-phase cells with enlarged and multiple nuclei (Fig. 7C).

**Fig. 7.**
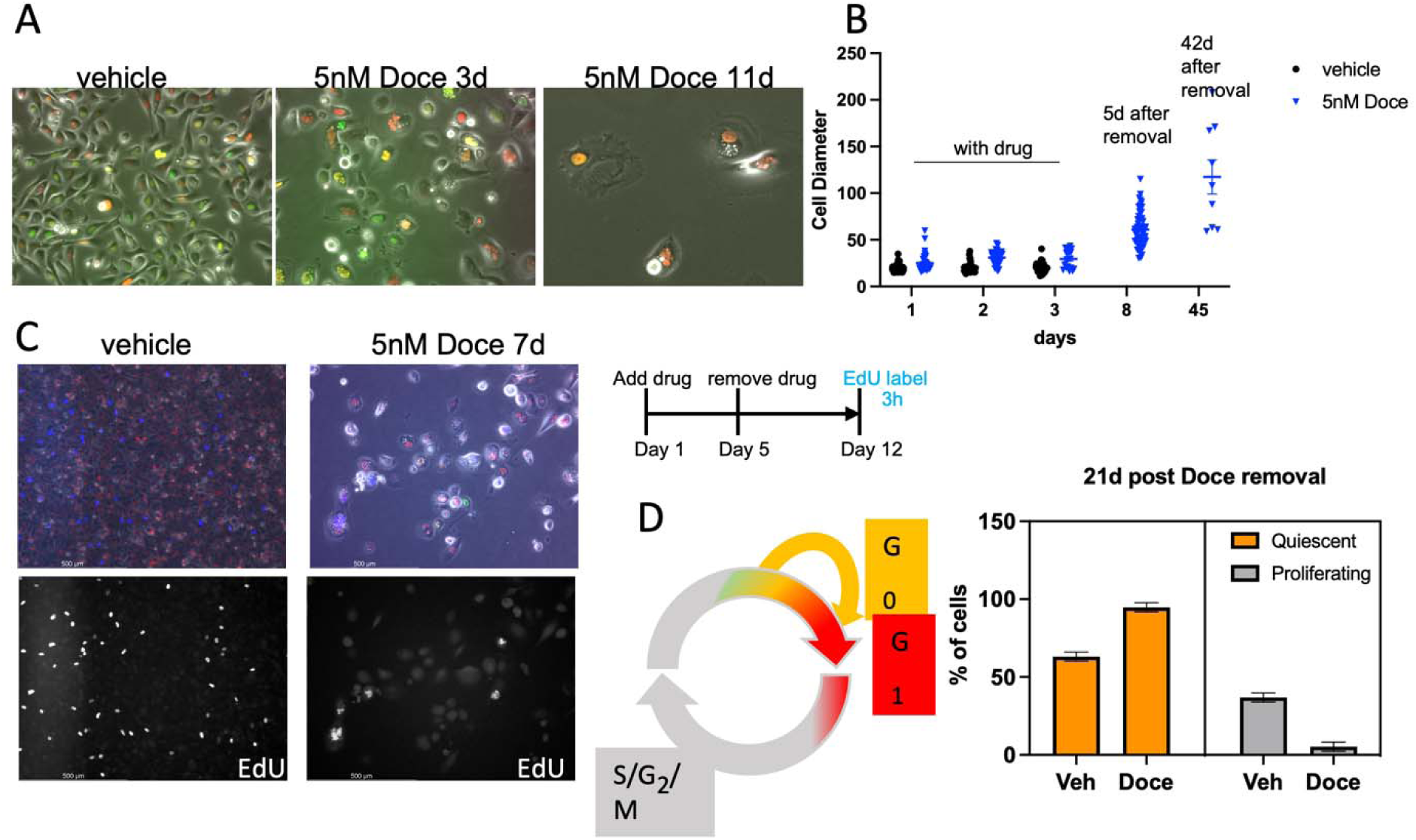
PC3 cells that survive Docetaxel treatment form large, polyploid cells. A. Cultured PC3 cells stably expressing cell cycle reporters p27K-Venus (G0) and Cdt1-Cherry (G0/G1), were treated with vehicle only (DMSO) or 5nM Docetaxel for 3 days, after which most PC3 cells die. B. After drug removal, long-term surviving PC3 cells continue to grow for over a month as shown by increasing cell diameter. C. EdU labeling reveals that PC3 cells that survive Docetaxel treatment endocycle. D. Using the p27K-Venus (G0) and Cdt1-Cherry (G0/G1) cell cycle reporters, we observe most long-term surviving polyaneuploid PC3 become quiescent. Note the vehicle treated PC3 cells at 21d are confluent and therefore also exhibit high levels of quiescence.

Our PC3 cell line is a clonal sub-line that stably expresses two cell cycle reporters, a p27-based Venus reporter that accumulates during the G0 phase of the cell cycle but is degraded during G1, S and G2/M (Oki et al., 2014), and a component of the FUCCI cell cycle reporter system, Cdt1-N-terminus (1-32) fused to mCherry which accumulates during G0 and G1 phases but is degraded during S-phase, G2 and M (Sakaue-Sawano et al., 2008). With these reporters, cell cycle arrest in G0 can be distinguished from cycling by live cell imaging as well as flow cytometry (Pulianmackal et al., 2021). We found that most surviving cells after docetaxel treatment become quiescent by 21d and remain in G0, with only a small fraction continuing to endocycle (Fig 7D).

The cell cycle arrest of surviving cells could be due to expression of cell cycle inhibitors in response to the endocycling state that results in increased genomic content. Indeed, we observed high levels of expression of the cell cycle inhibitor p21, along with high levels of phospho-ATM, both of which are induced by DNA damage and genetic instability. We also observed a small number of endocycling cells with high p16 expression, a cell cycle inhibitor which is normally epigenetically silenced in PC3 cells (Chi et al., 1997; Jarrard et al., 1997; Kondo et al., 2008), but may become expressed in a small subset of cells as cells enter a senescence-like state in response to sustained DNA damage and cell cycle arrest (Steiner et al., 2000) (Fig. 8A, B).

**Fig. 8.**
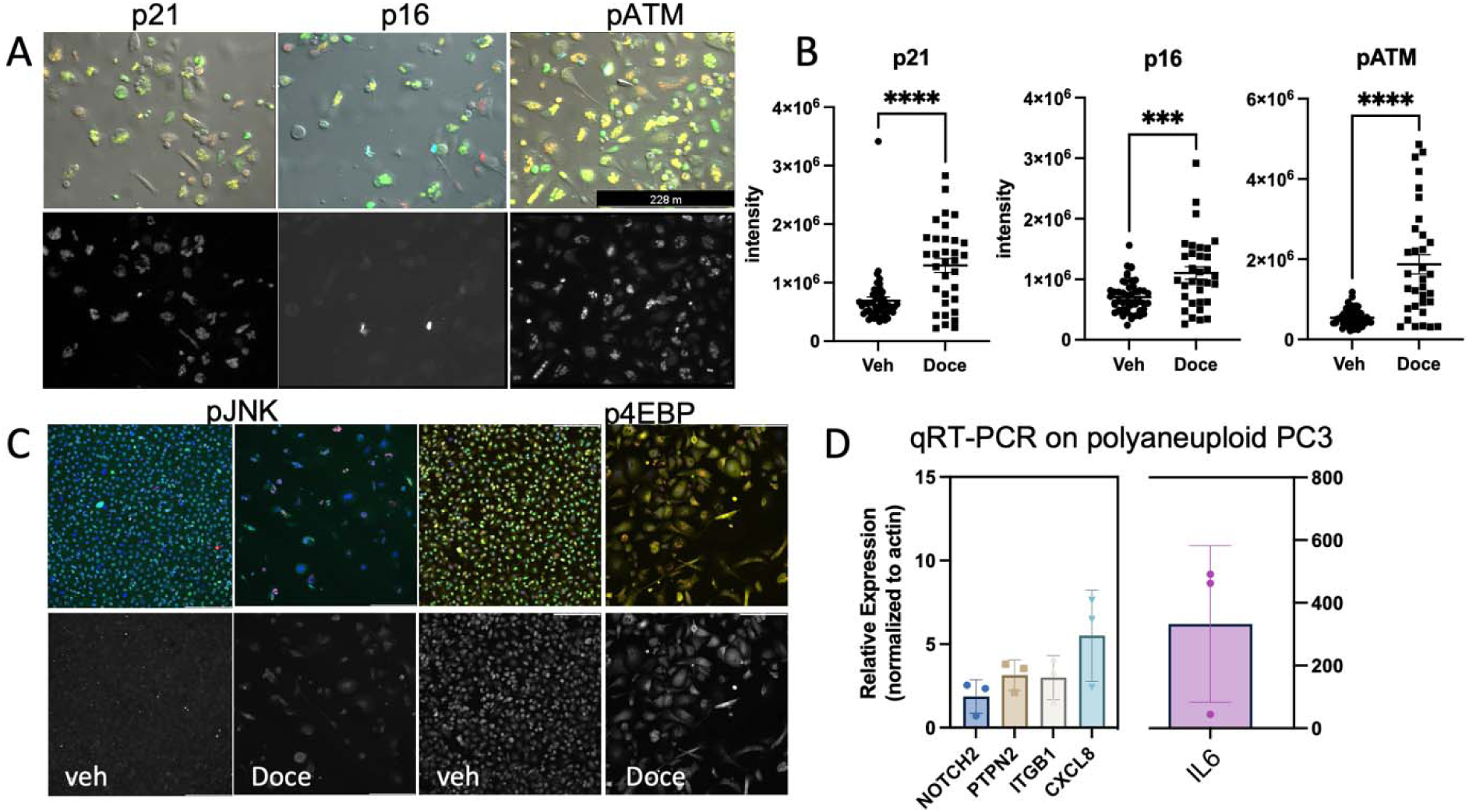
Surviving polyaneuploid PC3 cells express markers of cell cycle arrest, DNA damage and upregulate genes orthologous to those upregulated in hyperplastic *Drosophila* glands. A. Cultured PC3 cells stably expressing cell cycle reporters p27K-Venus (G0) and Cdt1-Cherry (G0/G1), were treated with vehicle only (DMSO) or 5nM Docetaxel for 3 days and 5 days after drug removal (7 days total), stained for markers of cell cycle arrest and DNA damage. B. PC3 polyaneuploid cells exhibit high levels of p21 and pATM expression, markers of DNA damage induced cell cycle arrest. PC3 polyaneuploid cells also occasionally exhibit high p16 indicating some may enter senescence. C. PC3 polyaneuploid cells 5 days after drug removal exhibit increased pJNK and p4EBP. D. qRT-PCR was performed on RNA isolated from surviving PC3 cells 10 days after drug removal for the indicated genes. Gene expression levels were normalized across samples using Beta-Actin with control RNA (from vehicle only treated cells) and set to 1 to indicate the fold-change in expression. Note that fold change for IL6 transcript is shown on the right y-axis, as avg. fold change exceeded 300.

We next examined whether the surviving polyaneuploid PC3 cells upregulate any of the signaling pathways we observed in hyperplastic postmitotic accessory gland cells. We observed upregulation of JNK signaling (via phosphorylation of JNK) and PI3K/TOR signaling (via phosphorylation of 4EBP) in surviving PC3 cells (Fig 8C). Consistent with the activation of these pathways, we also observed increased mRNA levels of human orthologs of several genes we see upregulated in hyperplastic *Drosophila* glands by qRT-PCR on PC3 polyaneuploid cells, including Notch (NOTCH2), Ptp61F (PTPN2), Integrin Beta (ITGB1), and human orthologs of the unpaired ligands, CXCL8 (encodes IL-8 protein) and IL6 (Fig 8D,E). These results suggest our hyperplastic postmitotic model in the *Drosophila* accessory gland may recapitulate specific features of human prostate cancer driven by cancer cells that survive by entering the endocycling state.

## Discussion

Here, we present an *in vivo* postmitotic and polyploid prostate cancer model in the *Drosophila* accessory gland. Our model displays many phenotypic hallmarks of tumorigenesis, such as disruption of polarity, apical invasion and basal extrusion, cellular and nuclear hyperplasia, as well as compromised tissue differentiation. What is unexpected about our *in vivo Drosophila* prostate hyperplasia model is that all these phenotypes could be activated in postmitotic cells, in the absence of proliferation. Our results reveal that postmitotic cells can activate a conserved network of tumorigenesis and wound-healing related genes, and likely engage in recently described feedback loops between a JNK-dependent senescent-like gene expression state and a JAK-STAT-induced pro-growth state that engages with the tissue microenvironment in an autocrine as well as paracrine manner (Jaiswal et al., 2023). Our results demonstrate that these feedback loops and gene expression programs are cell states that can be achieved entirely independent of cell proliferation, and can even, under rare conditions, result in neoplastic phenotypes without proliferation (Fig. 9). Finally, our investigations into prostate cancer cells that survive therapeutic insult by entering an endocycling state suggest these signaling states and gene expression changes are conserved in non-mitotic polyploid human prostate cancer cells.

**Fig. 9.**
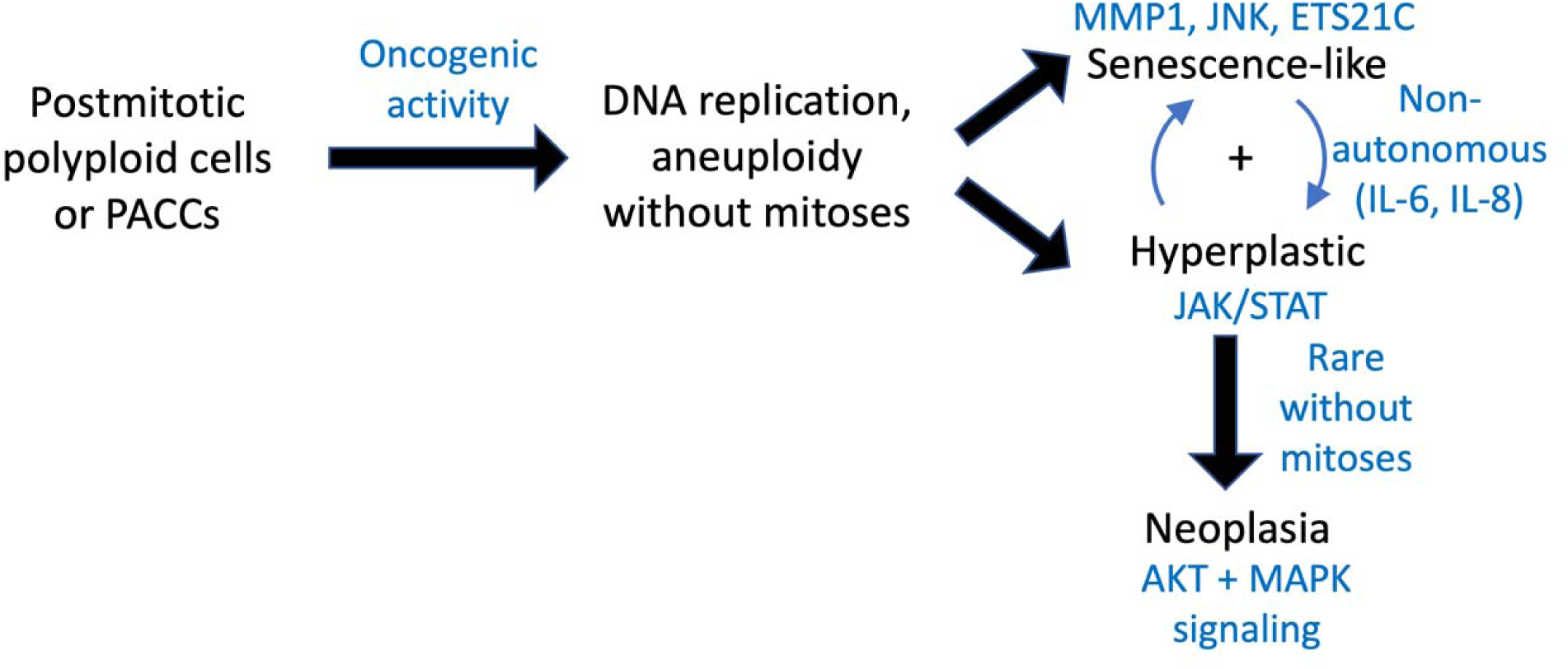
A model for non-proliferative pro-tumorigenic signaling in the adult *Drosophila* accessory gland and prostate polyaneuploid cancer cells.

### Non-proliferative polyploid cells in cancer

Not all cells in tumors proliferate. Polyploid cancer cells are frequently found in human cancerous tissues, with ∼37% of all human tumors reported in the Cancer Genome Atlas Pan-Cancer dataset showing evidence of polyploid DNA content (Bhathal et al., 1985; Liu, 2018). Studying polyploid progression and cancer phenotypes can be difficult. Polyploidy has been found to occur via several mechanisms, from increasing the number of nuclei, cytokinesis failure (Caldwell et al., 2007; Fujiwara et al., 2005; Meraldi et al., 2002), cell cannibalism by entosis (Augimeri et al., 2023; Krajcovic and Overholtzer, 2012), and cell fusion (Brodbeck and Anderson, 2009; Hosaka et al., 2004). Additionally, cells can become polypoid by increasing genomic content via endoreplication (Holland and Cleveland, 2009).

Polyploid giant cancer cells have been associated with resistance to cancer therapy in cell culture experiments (Barok et al., 2011; Kim et al., 2023; Lin et al., 2019; Liu, 2018), and can be induced by factors such as hypoxic conditions, chemotherapeutic drugs, and irradiation treatments *in vitro* (Fei et al., 2019; Kaur et al., 2015; Lanvin et al., 2013), which can result in increased migration and invasion capabilities, epithelial-mesenchymal transition (EMT) related protein expression, and MMP expression (Fei et al., 2019; Lv et al., 2014; Mallin et al., 2023; Wang et al., 2019; Zhou et al., 2022).

Our cell culture model for endocycling cancer cells is limited in that it is a monoculture. The chemotherapy induced endocycling prostate cancer cells exhibit JNK activation and gene expression changes similar to those observed in the *Drosophila* accessory gland after oncogenic activation. Most strikingly, we observe a very strong upregulation of IL6 and CXCL8 expression, consistent with senescence-like phenotypes in the polyaneuploid state (Niu et al., 2021; Pienta et al., 2022; Pienta et al., 2021). Interestingly, several Stats (STAT3, STAT5 A,B) are genetically mutated and lost in PC3 cells (Seim et al., 2017; Spiotto and Chung, 2000), suggesting the non-autonomous proliferative effects of the surviving large, polyaneuploid PC3 cells, perhaps through IL-6 and IL-8 release, should be examined in co-culture experiments with other prostate cell lines retaining the full Jak/STAT signaling pathway. This will also be important to examine in co-culture studies with immune cells and other cells of the prostate cancer microenvironment. PC3 cells have been shown to potentially signal via IL-6 protein in a paracrine manner (Lou et al., 2000). Future work should confirm the secretion of these paracrine factors by the cells in the large polyaneuploid state and examine their effects on other cells non-autonomously, to gain a deeper understanding of how cancer cells in this state impact the tumor microenvironment.

### The timing of oncogene activation and cell cycle exit

Activation of oncogenic Ras^V12^ expression during the proliferative phase of *Drosophila* accessory gland development was shown to frequently lead to neoplastic basal extrusions of cells that lack cell polarity and exhibit signaling consistent with a tumorigenic state (Rambur et al., 2020). In our study, we find that activation of oncogenic Ras^V12^ expression during the quiescent, mature phase of *Drosophila* accessory gland development leads to mild hyperplasia, but rarely progresses to neoplastic basal extrusions, despite high levels of MAPK and PI3K/Akt signaling. We attribute this to a loss of cell cycle and cell state plasticity after cell cycle exit and maturation in the adult accessory gland. The *Drosophila* accessory gland cells exit from the mitotic cell cycle ∼55-60h after puparium formation, but it is not known exactly when the accessory gland cells commit to their terminal cell fate. After the exit from the mitotic cell cycle, the gland undergoes additional endocycles, which appear to also occur in the Ras^V12^-induced neoplastic extrusions, suggesting cell fate is not totally compromised, although other aspects of epithelial differentiation, such as cell polarity, may be abrogated. We recently showed that the accessory gland main cells exit from the cell cycle ∼4-6 hours post eclosion in adults, after an additional round of endocycling, with only rare cells (<2% of the gland) continuing to endocycle (Box et al., 2024). As our genetic manipulations occur after exit from the endocycle, we speculate that the additional developmental time somehow reshapes and limits the accessory gland response to Ras^V12^ over-expression, which results in a milder phenotype with limited endocycling and growth despite robust MAPK and PI3K/Akt signaling. Studies comparing the gene expression networks induced by Ras^V12^ at different developmental times in this tissue may reveal the molecular basis for these differential effects of oncogenic activity.

Although neoplastic basal extrusions are rare in our study, we readily observe apical protrusions and invasions into the luminal space of the accessory gland (Fig. 1D-G). We also observe loss of proper cell-cell junctional protein localization, suggesting partial defects in cell polarity occur, even without total polarity loss as observed in basal extrusions. Epithelial polarity loss can lead to the aberrant localization of growth factor receptors, potentially creating feedback loops that promote tumorigenic signaling and impact cells non-autonomously. Additionally, the upregulation of MMP1, observed in our models, is indicative of JNK signaling, cellular stress, and active invasion into surrounding tissues. In cancer, even partial polarity loss can be associated with the acquisition of invasive capabilities, often marked by the enhanced expression of matrix metalloproteinases (MMPs).

In the proliferative developing fly eye, neoplastic-like conditions have been shown to generate polyploid cells that promote tumor growth. In these cells JNK signaling and Yki activity cooperate to promote growth while levels of the mitotic Cyclin, Cyclin B are suppressed, thereby preventing entry into mitosis leading to endoreplication (Cong et al., 2018). However, it is unclear in this developmental context whether the Cyclin B suppression occurs at a transcriptional or post-transcriptional level. While we see mRNA for several mitotic regulators increase in the hyperplastic-like endocycling accessory gland tissue (e.g. *aurA*, *aurB*, *cdk1*, *cycA, polo*), *cyclin B* transcript remains low, suggesting it may be transcriptionally silenced in the adult postmitotic accessory gland. An important difference in our model vs. the developing fly eye neoplastic-like tumors, is that the cells undergoing hyperplasia or neoplasia in the adult accessory gland are not already in a proliferative state. Further work will be needed to tease apart whether cycling vs. postmitotic endocycling cells in hyperplastic and neoplastic contexts engage different mechanisms to limit mitoses and induce polyploidy.

### JNK signaling, Ets21c induction and non-autonomous effects in the absence of proliferation

In the context of Myc+E2F, Myc+Ets21C, Ets21C, and E2F overexpression, evidence of excessive endoreplication and aneuploidy was observed, indicative of genomic instability. Aneuploidy can lead to proteotoxic stress due to imbalances in gene expression that result in JNK activation, ROS production, dysregulation of autophagy and, ultimately, hallmarks of tumorigenesis (Joy et al., 2024). Our RNAseq data suggest this also occurs in postmitotic cells of the accessory gland, as we also observe upregulation of genes involved in DNA damage, genetic instability, response to ROS and protein misfolding. Our gene expression profiles indicate strong upregulation of JNK signaling, with activation of known targets such as *puckered*, *Ets21C* and *MMP1*. In addition, recent work has revealed non-autonomous signaling induced by a senescence-like program downstream of JNK signaling (Jaiswal et al., 2023) that can also be activated in non-dividing polyploid cells under pro-growth conditions (Huang et al., 2024).

In *Drosophila*, Ets21C is upregulated in response to JNK signaling (Kulshammer et al., 2015; Toggweiler et al., 2016) as well as inactivation of the transcriptional repressor *capicua* (Jin et al., 2015). Here we show JNK signaling is increased in prostate cancer polyaneuploid cells and human Capicua (Cic) has been shown to be a tumor suppressor in prostate cancer (Gupta et al., 2022; Seim et al., 2017), suggesting the *Drosophila* accessory gland recapitulates key signaling features of the human cancer, despite the postmitotic state of the fly prostate-like cells. Capicua function is downregulated by Ras/Map Kinase signaling, through degradation, nuclear export, or both (Astigarraga et al., 2007; Grimm et al., 2012; Jin et al., 2015; Roch et al., 2002; Tseng et al., 2007). Although we do not observe MAPK activity via phosphorylated ERK in most of our accessory gland hyperplasias, we do find high phosphorylated ERK in hyperplasias exhibiting basal extrusion. The high JNK signaling induced in accessory gland hyperplasias may promotes Ets21c activation, while the additional MAPK activation observed in basal extrusions may further elevate Ets21C via Cic inactivation to drive cell cycle targets to induce endocycling. Both Ets21C and Cic directly impact cell cycle targets involved in endoreplication and DNA synthesis, with Cic acting as a repressor and Ets21C acting as an activator (Jin et al., 2015; Zhang et al., 2022). These pathways converge on genes that promote endocycling activated by Myc and E2F leading to the excessive endocycling and nuclear anaplasia we observe in Myc+E2F conditions. The upregulation of *Drosophila* Ets21C in the accessory gland in response to oncogenic signaling may be particularly significant, considering its homology to the human ERG gene and other ETS transcription factors, known for their involvement in prostate cancer (Adamo and Ladomery, 2016; Nicholas et al., 2019; Qian et al., 2022; Tomlins et al., 2005). Our data suggest Ets21C plays roles in promoting endocycling and cell growth both cell autonomously and non-autonomously, consistent with studies revealing paracrine targets of Ets21C in proliferative contexts (Mundorf et al., 2019; Toggweiler et al., 2016). Our *Drosophila* model offers an opportunity to further examine the role of Ets21C in promoting tumorigenic signaling in non-mitotic cells and may suggest a role for ERG in polyploid cells of prostate tumors.

### A common tumor gene expression profile occurs even in the absence of proliferation

Our transcriptomic analysis of the response to E2F+Myc and E2F + Yki* in the *Drosophila* accessory gland, revealed gene expression changes that overlap significantly (10.97 log_2_ - fold greater than expected by chance) with a common wound healing and tumorigenesis gene expression program (Floc’hlay et al., 2023). Interestingly, our results suggest the shared commonality in the enhancer gene regulatory networks of tumor behavior and the response to wounding, can also be engaged even in the absence of cellular proliferation (MacCarthy-Morrogh and Martin, 2020). This work is the beginning of disentangling the contribution of cell cycle plasticity to tumorigenic gene expression networks, and further studies in tissues with variant cell cycles may reveal additional aspects of tumorigenic programs that are proliferation independent.

## Acknowledgements

We thank Dr. Mirka Uhlirova for kindly providing the UAS-Ets21C flies. This work was supported by The Prostate Cancer Foundation through a 2016 Challenge Grant (16CHAL05) to LB, KP and FC and an American Cancer Society Scholar Award (RSG-15-161-01-DDC) to LB, NIH/NIGMS R01GM127367 to LB, an NIH/NIGMS R35GM149273 to LB, an NIH/NCI P01CA093900 to SA and KP and an CDMRP/PCRP Idea Development Award (W81XWH-20-10353) to SRA and LB. SJC was supported by a Rackham Merit Fellowship, University of Michigan. We thank Dr. Russell Taichman for extensive advice and help during the development and execution of this project.

## Conflicts of Interest

Kenneth Pienta discloses that he is a consultant to Cue Biopharma Inc., an equity holder in PEEL therapeutics, an equity holder in Keystone Biopharma Inc, and an equity holder in Kreftect, Inc. Sarah Amend discloses that she is an equity holder in Keystone Biopharma Inc.

## Material and Methods

### Staining and Fixation

*Drosophila* samples and dissected accessory glands were fixed in a 4% paraformaldehyde/1× PBS solution for 30 minutes, followed by three 10 min. washes in 1× PBS with 0.1% Triton X-100 (PBST). The samples were then permeabilized with 0.5% Triton X in PBS for 30 minutes and blocked in a solution of PBS, 0.1% Triton X-100, and 1% bovine serum albumin (BSA) for 10 minutes. Primary antibody incubation was conducted in PAT at 4°C overnight. This was followed by three washes in PBST and incubation in secondary antibodies conjugated with the required fluorophores, mixed in PBS with 0.3% Triton X-100, 0.1% BSA, and 2% normal goat serum (PBT-X+2%NGS) at room temperature for 4 hours or overnight at 4°C. For nuclear staining, DAPI was applied 1:1000 in PBST for 10 min, followed by three rinses of PBST, for 10 min each. The samples were mounted on glass slides using Vectashield mounting medium. Imaging was performed using a Leica SP5 confocal microscope.

Primary antibodies used in this study include: Phospho Histone H3 Ser10 Mitosis Marker mouse 1:500 (Millipore 06-570), MMP1 catalytic domain mouse 1: 10 (DSHB 5H7B11-s, 3B8D12-s, 3A6B4-s), Dcp1 Cleaved *Drosophila* rabbit 1:1000 (Cell Signaling 9578s Asp216), Integrin Beta PS mouse 1:100 (DSHB CF.611), Discs Large mouse 1:500 (DSHB 4F3), MPM-2 Mitotic Protein Monoclonal mouse 1:100 (Millipore 05-368), Lamin mouse 1:300 (DSHB ADL67.10), DP-MAPK 1:500 (Cell Signaling 4370s), Fas III 1:500 (DSBH 7G10), Coracle-heavy 1:100 (DSHB C615.16-s), P-Akt 1:200 (Cell Signaling 4054s), P-4E-BP 1:200 (Cell Signaling 2855S). Secondary antibodies used in this study are: Alexa Fluor 568 goat anti-mouse 1:2000 (Invitrogen A11031) or Alexa Fluor 568 goat anti-rabbit 1:2000 (Invitrogen A-11011), Alexa Fluor 594 F(ab’)2 goat anti-mouse 1:2000 (Invitrogen A11020), Alexa Fluor 594 F(ab’)2 goat anti-rabbit 1:2000 (Invitrogen A11072).

### EdU labeling

EdU labeling was performed as instructed by manufactures protocol, ‘Click-iT™ Plus EdU Cell Proliferation Kit for Imaging, Alexa Fluor™ 555 dye’ (Thermo Fisher C10638). For EdU assays male flies were exposed to a 37 °C water bath for 20 minutes on day of eclosion. Animals were housed in a 25°C incubator and aged. Male flies were aged for 6 days post eclosion before being fed 1 mM EdU in 10% sucrose with blue food coloring for the four days. A mixture of EdU and sucrose was applied to Whatman paper, which was then placed inside empty vials. This setup was refreshed every two days to prevent contamination.

### Fly rearing and mating

Fly Stocks

*w/w; UAS-P35/Cyo-GFP; Act>CD2>gal4, UAS-GFP(nls)/TM3-Ser-GFP* (UAS-P35 BL#5072, act>CD2>Gal4 from BL#4780)

*y,w,hs-flp^12^; +; UAS-yki* (yki from BL#28819)

*y,w,hs-flp^12^; UAS-E2F, UAS-DP/Cyo-GFP; yki\** (UAS E2F1,UAS DP from Neufeld et al 1998, Constitutively active UAS-Yki* from BL#28817)

*y,w,hs-flp^12^; UAS-E2F, UAS-DP/Cyo-GFP; UAS-dMyc42/TM6B* : (UAS E2F1,UAS DP

from Neufeld et al 1998, dMyc from BL#9675)

*y,w,hs-flp^12^; + ;UAS E2F1, UAS-Dp/TM6B* (BL#4770)

*y,w,hs-flp^12^; +; +*

*y,w,hs-flp^12^; UAS-Rasv12/ CyO-GFP; +* (BL#5788 or BL#64196)

*y,w,hs-flp^12^; +; UAS-Ykis111,168,250/TM6B* (*BL#28817)*

*y,w,hs-flp^12^; +; UAS-dMyc/TM6B* (dMyc from BL#9675)

*w/w; UAS-Ets21C/Cyo-GFP; Act>CD2>gal4, UAS-GFP(nls)/TM6B* (Ets21C from Mirka Uhlirova et al 2015, act*>CD2>Gal4 from BL#4780)*

*w/w; UAS-Ets21C-RNAi/Cyo-GFP; Act>CD2>gal4, UAS-GFP(nls)/TM6B* (*Ets21C-RNAi* VDRC 318056, *act>CD2>Gal4 from BL#4780)*

For antibody staining male flies 1-3 days after eclosion had transgene expression induced by exposing animals to a 37 °C water bath for 20 minutes. Animals were housed at RT and aged for various timepoints.

### Imaging/Measurements/DAPI quantifications

Imaging was conducted using a Leica SP5 confocal microscope at various magnifications, with a Z-section thickness of 1 μm. Maximum intensity projections were created using LAS AF software. Image quantification was carried out with FIJI software. For this process, regions of interest (ROIs) of similar size were manually outlined based on nuclear (DAPI) staining or antibody staining. To normalize the area for quantification, the integrated density was adjusted by subtracting the background measured in ROIs outside of the tissue. Data for nuclear intensity and line-scans are available in Supplement.

### Flow Cytometry

We followed the accessory gland nuclear isolation and flow cytometry from (Box et al., 2024) for ovaries and accessory glands. Vybrant DyeCycle Violet stain (Invitrogen) at 1:1000 was used for nuclear staining. Samples were lightly vortexed just prior to running on an Attune Flow Cytometer.

### RNA Bulk Seq

Male flies, 10 days post-eclosion, were selected for dissection after visually confirming high transgene/GFP expression. Their accessory glands were extracted and immediately placed in TRIzol (Invitrogen 15596026) for RNA preservation. Following the manufacturer’s instructions, we isolated RNA from these samples. The prepared RNA samples were then sent to the University of Michigan’s Sequencing Core for further analysis.

At the sequencing facility, we employed PolyA selection to create barcoded libraries for each sample. The integrity and quality of these libraries were verified using a Bioanalyzer and qPCR. We conducted sequencing on the NovaSeq SP 100 cycle platform, ensuring high-read quality through FastQC assessments.

For data analysis, sequences were aligned using the Sleuth/Kallisto pipeline, with reference to the *Drosophila* melanogaster genes based on the Ensembl BDGP6.28 database. The complete dataset will be made publicly available on the Gene Expression Omnibus (GEO) database. Differentially expressed genes were identified based on a log2 fold change threshold of ±1 (equivalent to a 2-fold change) and an adjusted P-value of less than 0.05, unless otherwise stated see supplemental Fig. 10-11.

### Cell culture

PC3 cells expressing G0-Venus and G1-Cherry fluorescent reporters were cultured as described previously (Pulianmackal et al., 2021). For induction of surviving endocycling cell state, cells were seeded into 12-well plates or chamber slides at a density of 1-5 million cells/mL and treated with vehicle (DMSO) or 5nM Docetaxel (Cayman Chemical) in DMSO for 72h. Media containing vehicle or Docetaxel was removed, adhering cells were washed with 1X PBS and fresh media without DMSO or Docetaxel was added, and cells were cultured for the indicated number of days post drug removal. For EdU labeling, 10µM EdU was added to the media for 20h prior to cell fixation in 4% PFA/PBS. EdU incorporation was assayed using an Alexa-Fluor 647 Click-IT EdU detection kit.

### qRT-PCR

For qRT-PCR assays, total RNA was isolated from vehicle treated PC3 or docetaxel treated PC3 cells at 10 days after drug removal. Reverse transcription with polyT and random primers was performed to generate cDNA, which was subsequently used for qRT-PCR. Primer sequences for qRT-PCR assays are provided in the supplement.

### Statistical Analysis and Visualization

Our data were analyzed, statistically tested, and visualized using a suite of Python packages including Pandas, NumPy, Matplotlib, Matplotlib_venn, Seaborn, Networkx, Scipy, and Statsmodels.api. Additionally, we employed Attune and LAS AF software, along with the R package Sleuth, for assessing differential expression and calculating fold changes.

For the EdU comparisons (Fig 5), we applied the two-sided Wilcoxon–Mann–Whitney test using SciPy to compare experimental genotypes against the control condition. In analyzing nuclear intensity (Fig. 6), we leveraged statsmodels.api to conduct Mixed Linear Model Regression, comparing our experimental genotypes with the control. This dataset included 200 observations across 4 biological replicates per condition, with the Mixed Linear Model Regression offering a robust framework to account for within-replicate variability, unlike the Wilcoxon–Mann–Whitney test.

Line-scan data (Fig. 2) generated from Fiji were analyzed using NumPy for quartile range calculations and Matplotlib for visualization. We employed the Wilcoxon–Mann– Whitney test for Violin plots to compare GFP-positive nuclear intensity against non-GFP nuclear intensity (Fig. 6). Venn diagrams were created with Matplotlib and Matplotlib_venn, (Fig. 4) while the online tool InteractiVenn was used for generating supplementary venn diagrams (Fig. 12).

For constructing node graphs (Fig. 4), we used Networkx and Matplotlib, focusing on genes with an adjusted p-value of 0.05 or lower and a fold change of ±1 or greater. GO-Term analysis of differentially expressed genes, adhering to the same thresholds, was conducted using the NIH’s DAVID for Biological Process categorization. Only GO-terms meeting these significance and fold change criteria were selected.

When creating heatmaps, we utilized gene ontology datasets from Amigo 2, with differential gene expression thresholds for adjusted p-value and fold change as outlined for each genotype compared against the Amigo 2 datasets in the supplementary materials Fig. 6-9.

### Fold enrichment and Hypergeometric distribution

Fold enrichment calculations is a ratio comparing observed frequency in the subset to expected frequency if the distribution were random. Hypergeometric calculations, use the hypergeometric distribution to calculate the probability of observing overlap between two or more sets, considering the sizes of the sets and the total population. The hypergeometric distribution probability calculation is a statistical method used in bioinformatics to determine whether a set of genes (or other elements) is overrepresented in a subset of a larger set. Additional details are provided in the supplement.

## Notes

### Summary of Updates

Funding statement revised to conform to NIH preferred format.

