## Supplement for "Oncogenic signaling in the adult *Drosophila* prostate-like accessory gland leads to activation of a conserved pro-tumorigenic program, in the absence of proliferation" (from another version of this preprint: version 1, 2024-05-12)

**Supplement to Church et al.**

**Supplemental Figures**

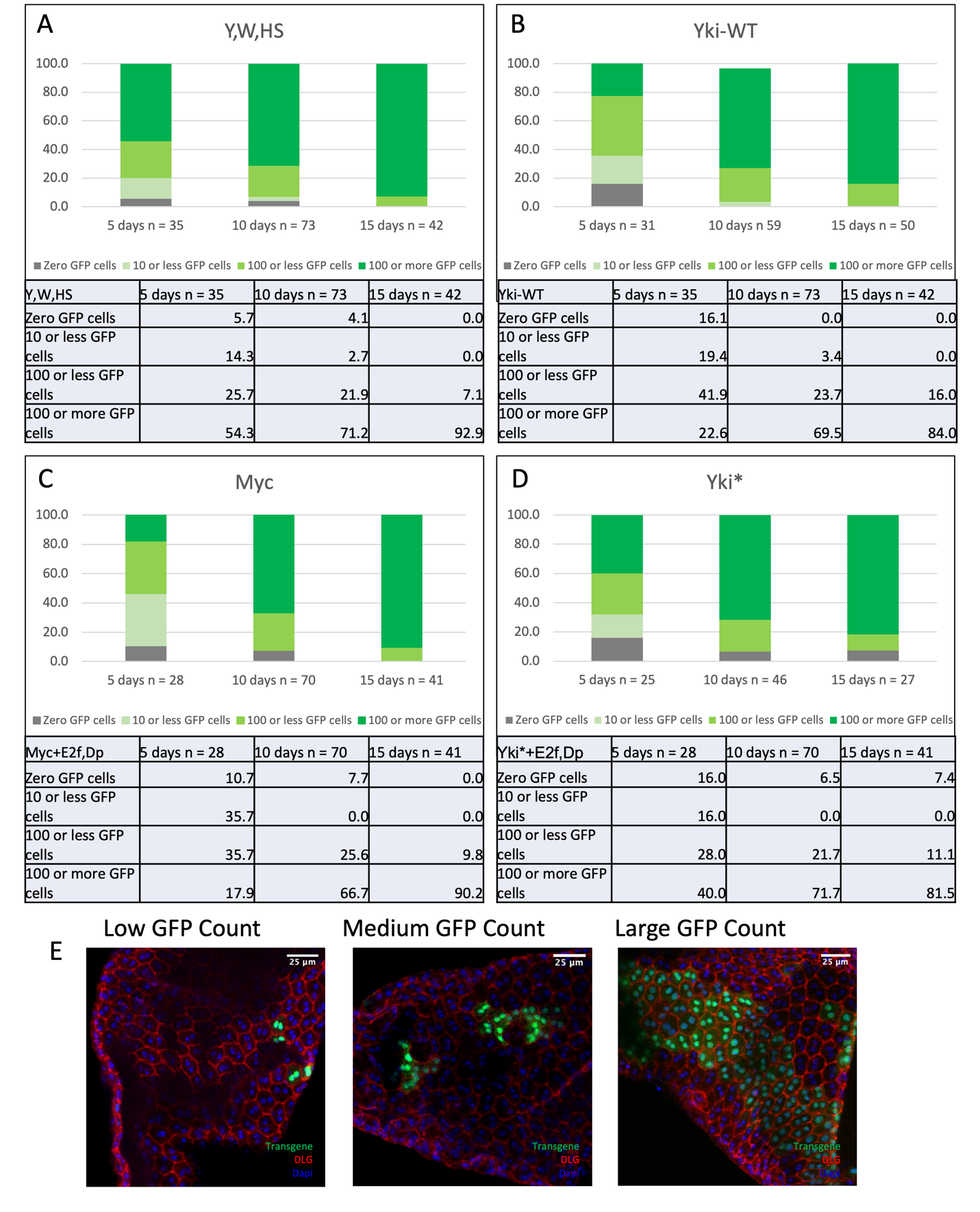

**Supp. Fig 1. Heat-shock parameters and Gal4/UAS-GFP expression for adult accessory gland.** Figures A-D, show samples that were heat-shocked for 20 minutes in a 37ºC water bath, then aged for 5, 10, and 15 days. After aging flies were dissected and the number of GFP positive cells were scored. Figure E shows representative images of the GFP scoring system.

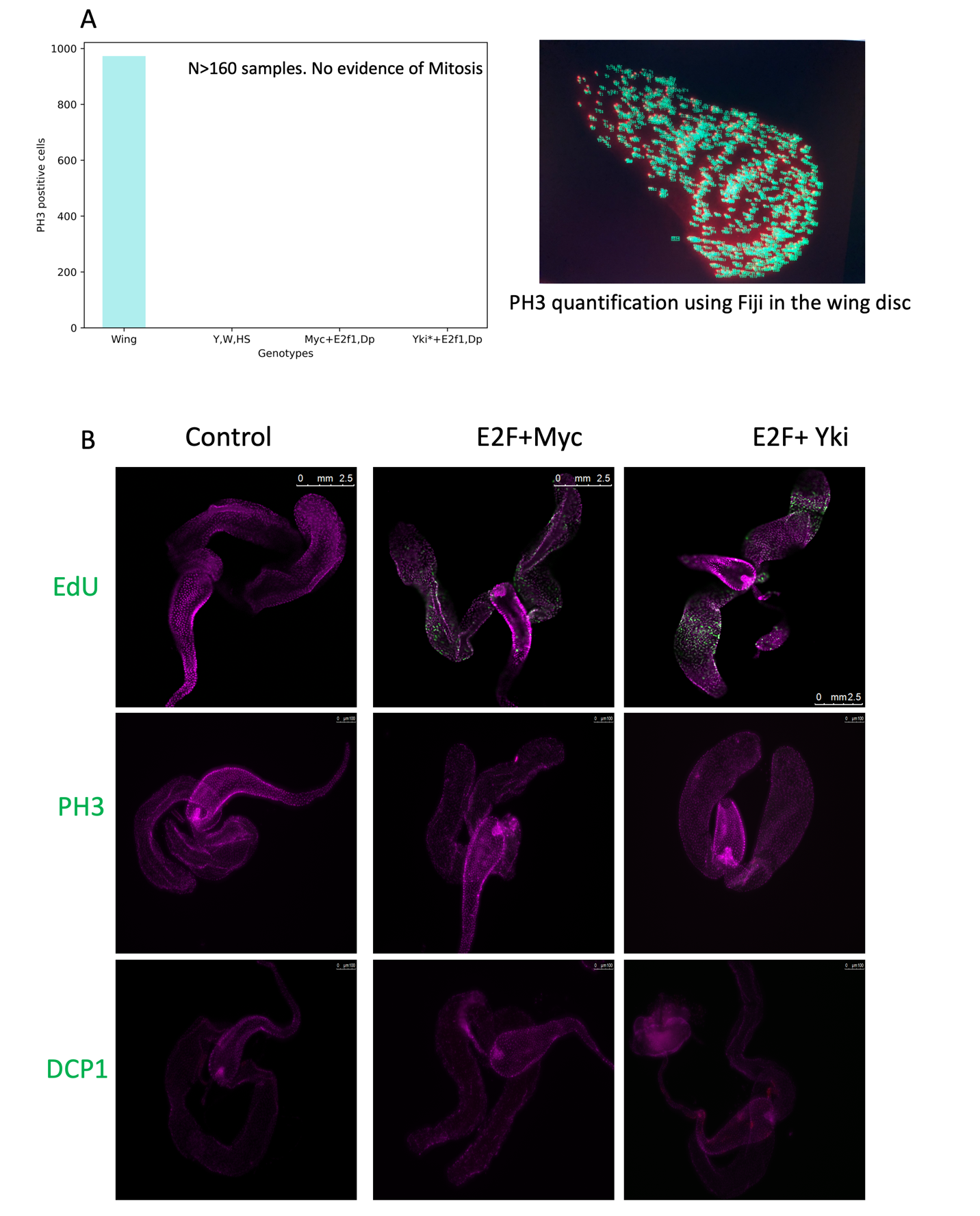

**Supp. Fig. 2 Mitotic and EdU labeling in accessory gland.**

A) Larval wing discs were co-stained in the tube with each accessory gland sample to ensure the mitotic marker labeling was working. PH3 quantifications show no PH3 positivity in glands but extensive mitoses in wings. B) Additional examples of EdU labeling for the indicated genotypes. E2F+Myc and E2F+Yki glands are negative for PH3 despite extensive endocycling and EdU labeling. We also observe no evidence of cell death, as assayed by anti-DCP1 antibody.

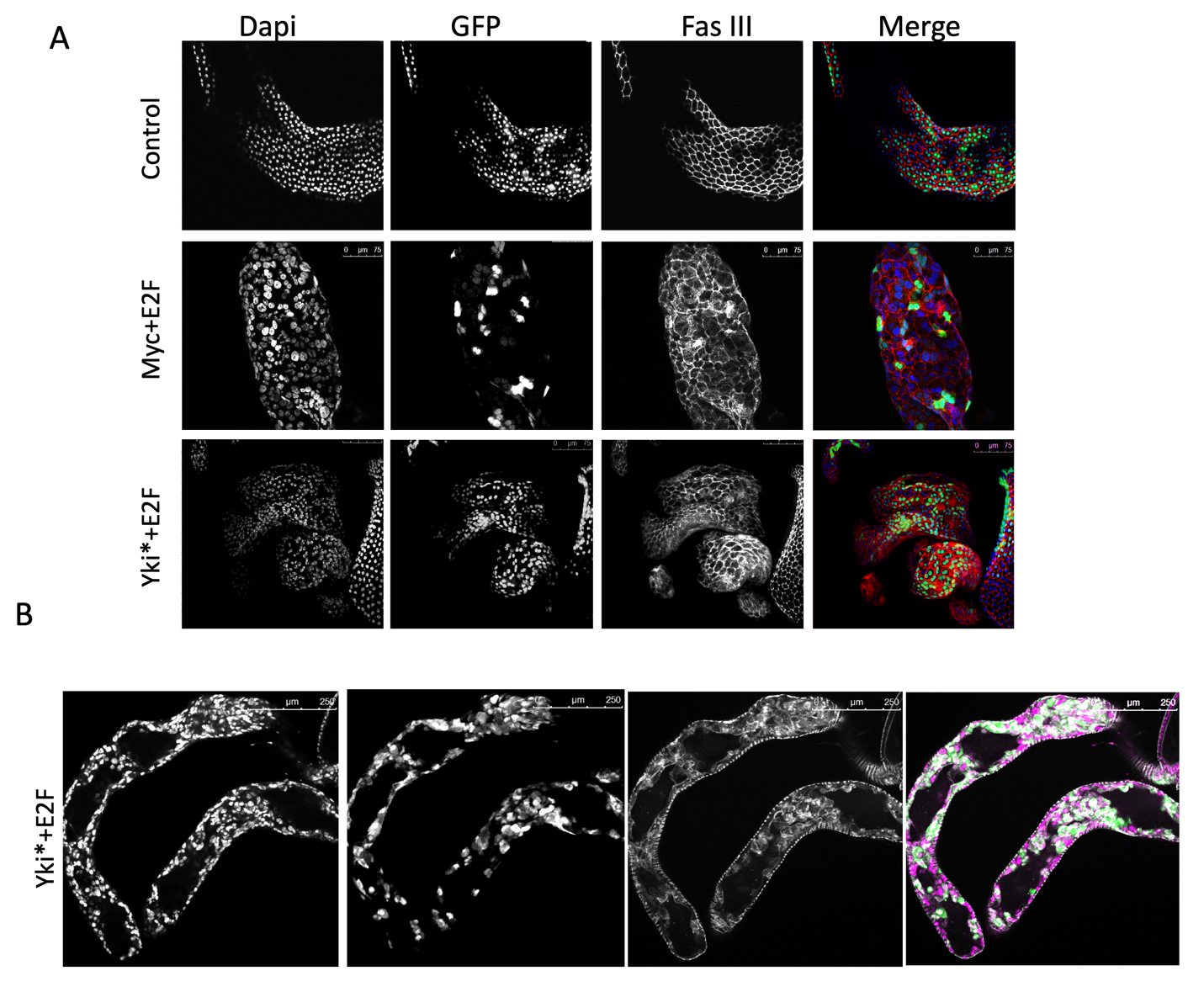

**Supp. Fig. 3 Septate junction protein Fas3 mis-localization and hyperplasia in accessory glands.**

A) Control (expressing GFP and P35), Myc+E2F and Yki*+E2F glands ten days after heat-shock. Myc+E2F and Yki*+E2F glands show Fas III mis localization. Sample sizes: Control N = 5, Myc+E2f N = 4, Yki*+E2f N = 6. B) Z-sections through the lumen of glands expressing Yki*+E2F fifteen days after heat-shock. Luminal invasion and hypertrophy are observed.

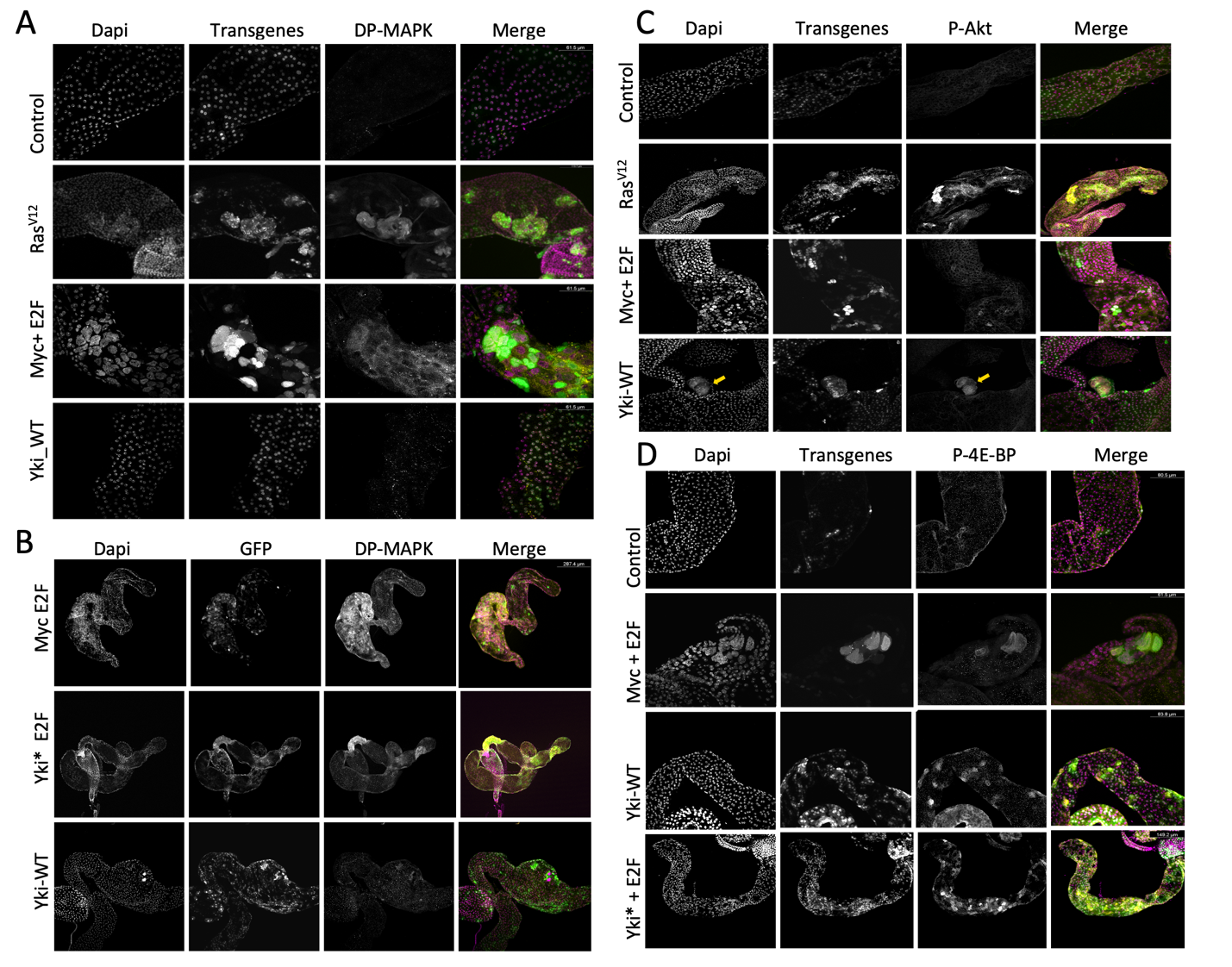

**Supp. Fig. 4 Additional examples of MAPK, PI3K/Akt signaling.**

A) dp-MAPK staining of samples at 5-20d post transgene induction. Most Myc+E2F and Yki^WT^ or E2F+Yki* samples are negative for dp-MAPK (note gain was increased to show even low level of staining in the Myc+E2F sample), while Ras^V12^ samples are always positive. B) Extreme examples of Myc+E2F-induced hyperplasia or examples of E2F+Yki* with extreme morphological defects (eg. misshapen gland structure) are dp-MAPK positive, suggesting when the ECM is disrupted, MAPK signaling is activated. Sample sizes: Myc+ E2f, 15 days N =6, Yki*+E2f 20days N = 10, and Yki-WT 20days N = 11, Ras^V12^ 5days N = 6, Myc+ E2F 10-days N =6, Yki-^WT^ 10days N = 14, and control (GFP+P35 only) 15days N = 8. C) P-Akt staining of samples at 10 days. Note only a neoplastic-like mass in Yki^WT^ and Ras^V12^samples are strongly positive. Control N = 12, RasV12 N = 2, Myc+ E2f, Dp N = 11, Yki*+E2f, Dp N = 9, and Yki-WT N = 14. D) PI3k/mTORC1 signaling assessed by p4EBP. All samples except for controls (GFP+P35 alone) are positive with or without hyperplastic growth at 10days. Note that some samples overlap with those shown in Fig 3.

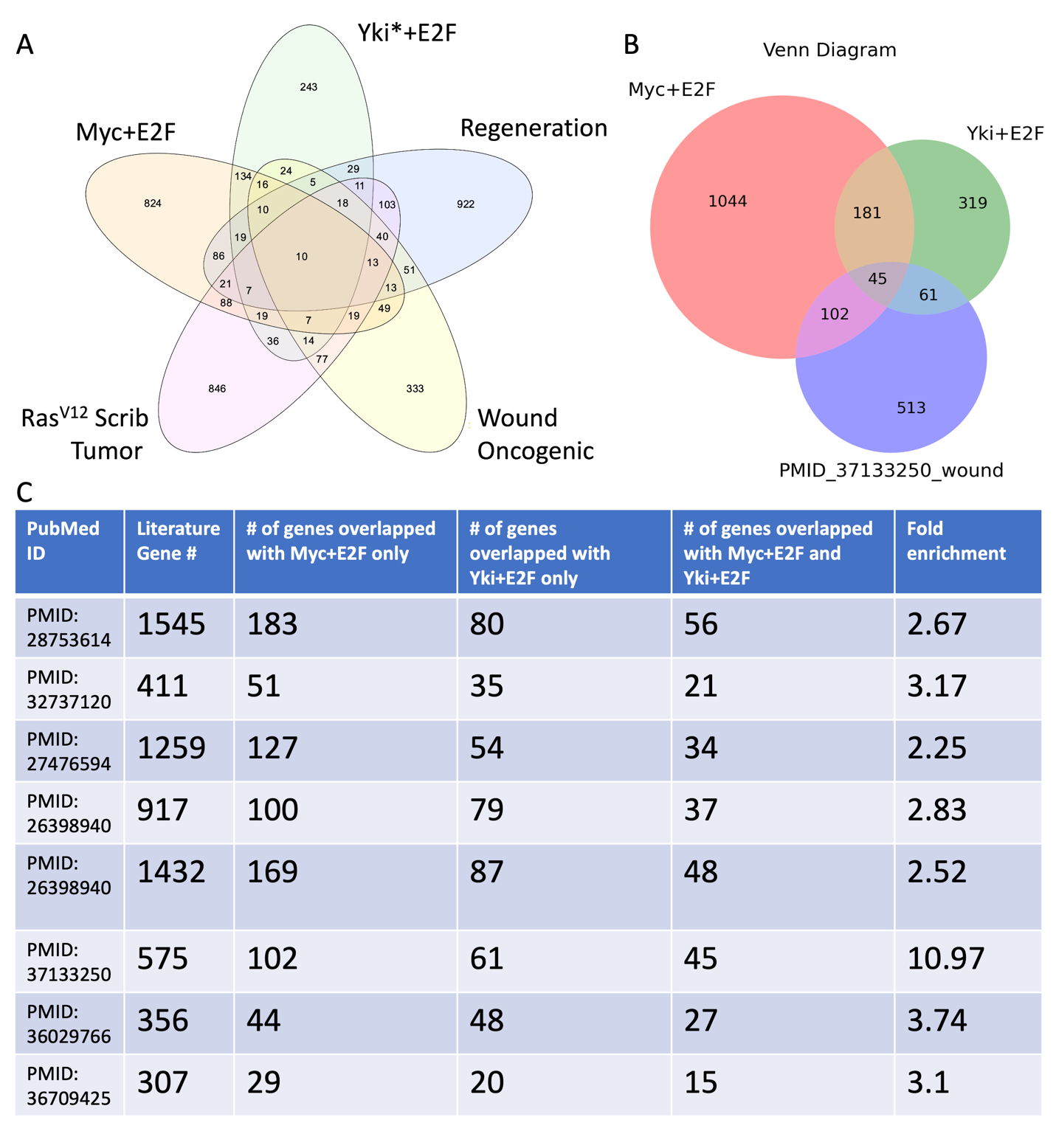

**Supp. Fig. 5 Overlap of wounding and tumorigenic gene expression signatures**

A) Five-way Venn diagram to show overlap of Myc+E2F, Yki*+E2F, PMID: 28753614, PMID: 37133250, and PMID: 26398940 datasets using the online tool Interactivenn.net. Note this venn diagram is not to scale. B) Three-way Venn diagram showing overlap of our datasets with a wounding gene expression response. C) Details of overlap between our data and the indicated published studies. The fold-enrichment of overlap, beyond that expected by chance (with chance = 1) using the hypergeometric calculation.

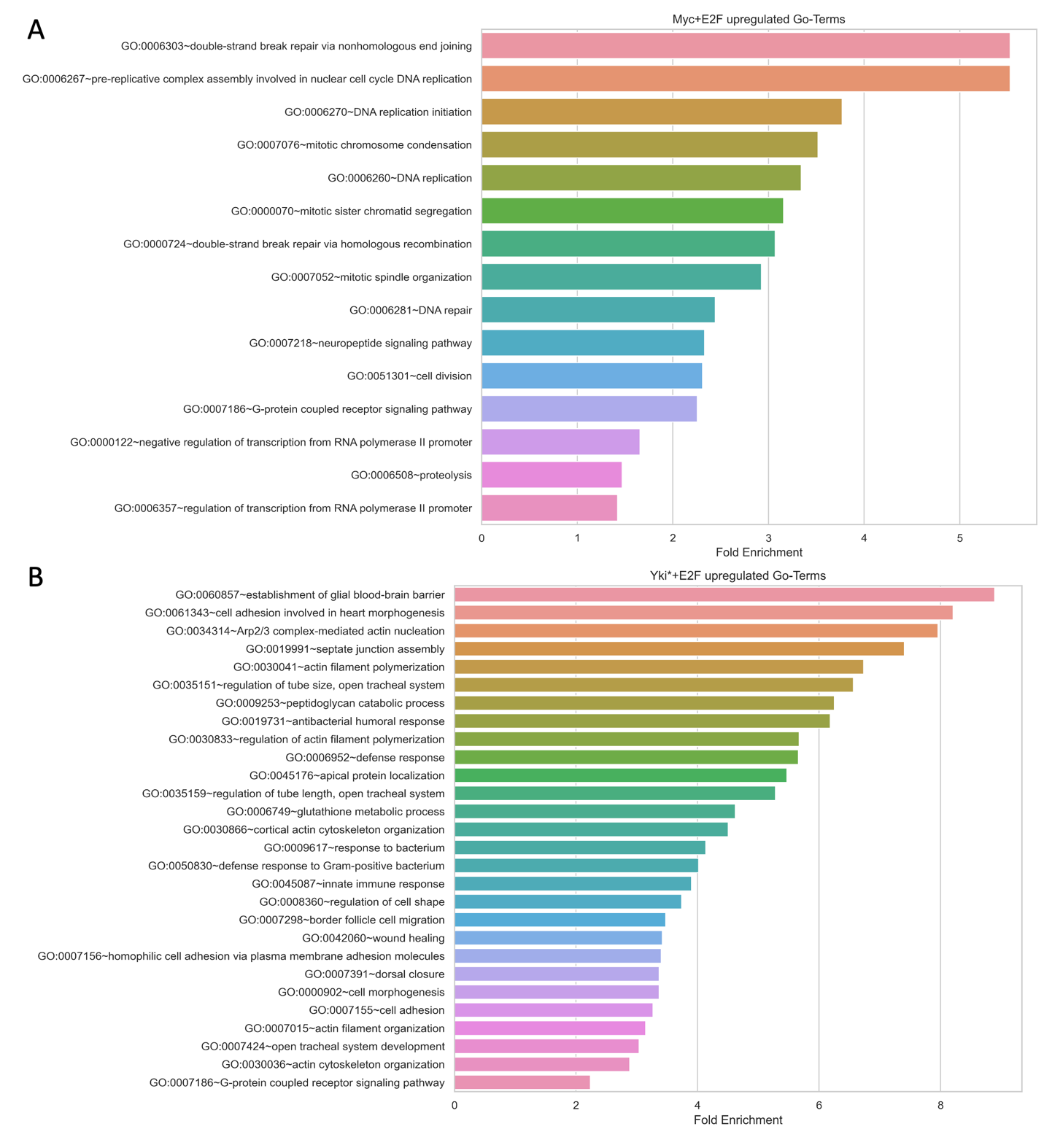

**Supp. Fig. 6 GO Term analysis of genes upregulated in hyperplastic glands.**

GO term enrichment analysis was performed using DAVID and multiple redundant terms were collapsed using ReviGO. Analysis was limited to genes with an adj p-value of 0.05 and log_2_-based fold changes of 1 or greater.

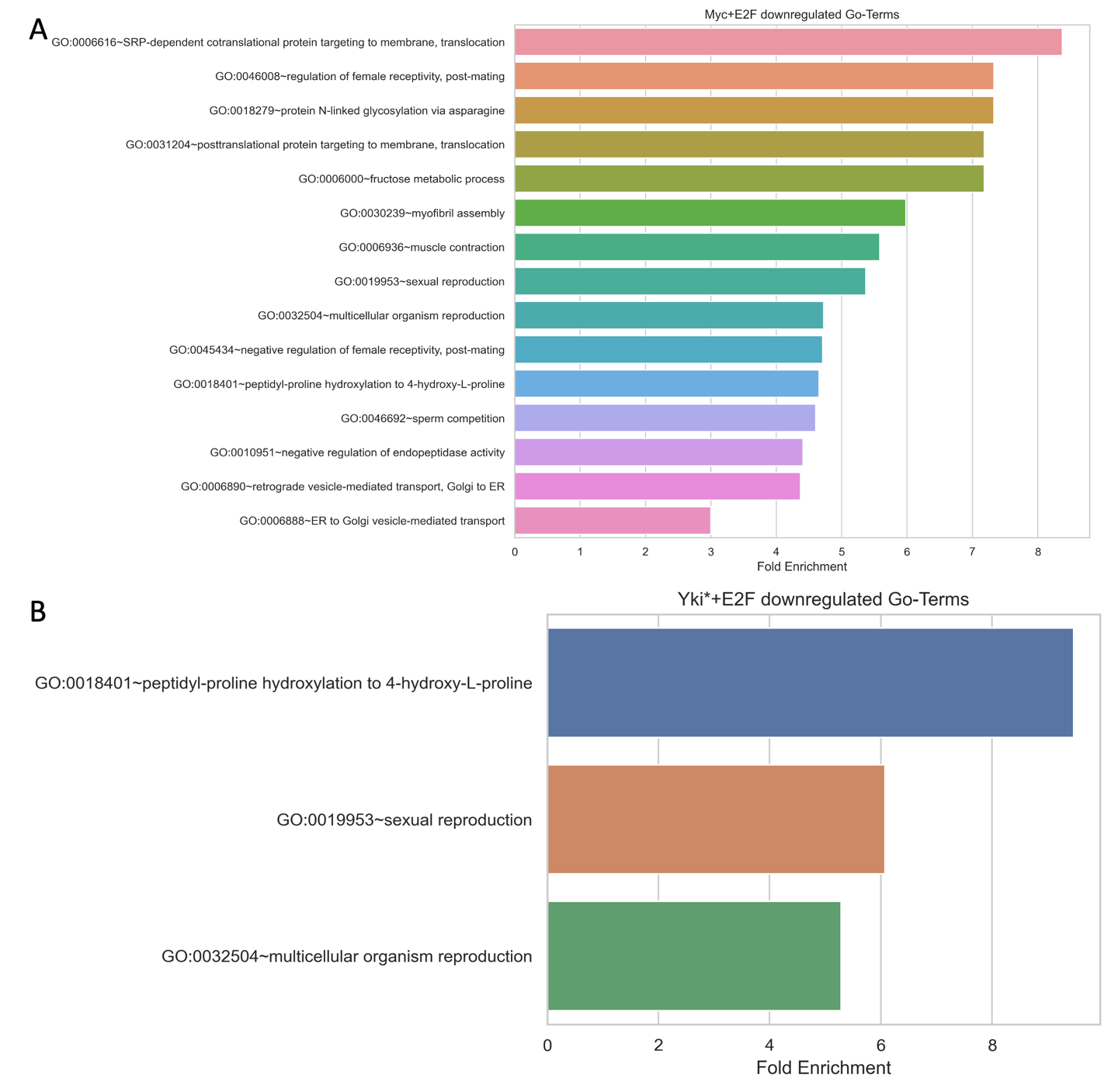

**Supp. Fig. 7 GO Term analysis of genes downregulated in hyperplastic glands.**

GO term enrichment analysis was performed using DAVID and multiple redundant terms were collapsed using ReviGO. Analysis was limited to genes with an adj p-value of 0.05 and log_2_-based fold changes of -1 or greater.

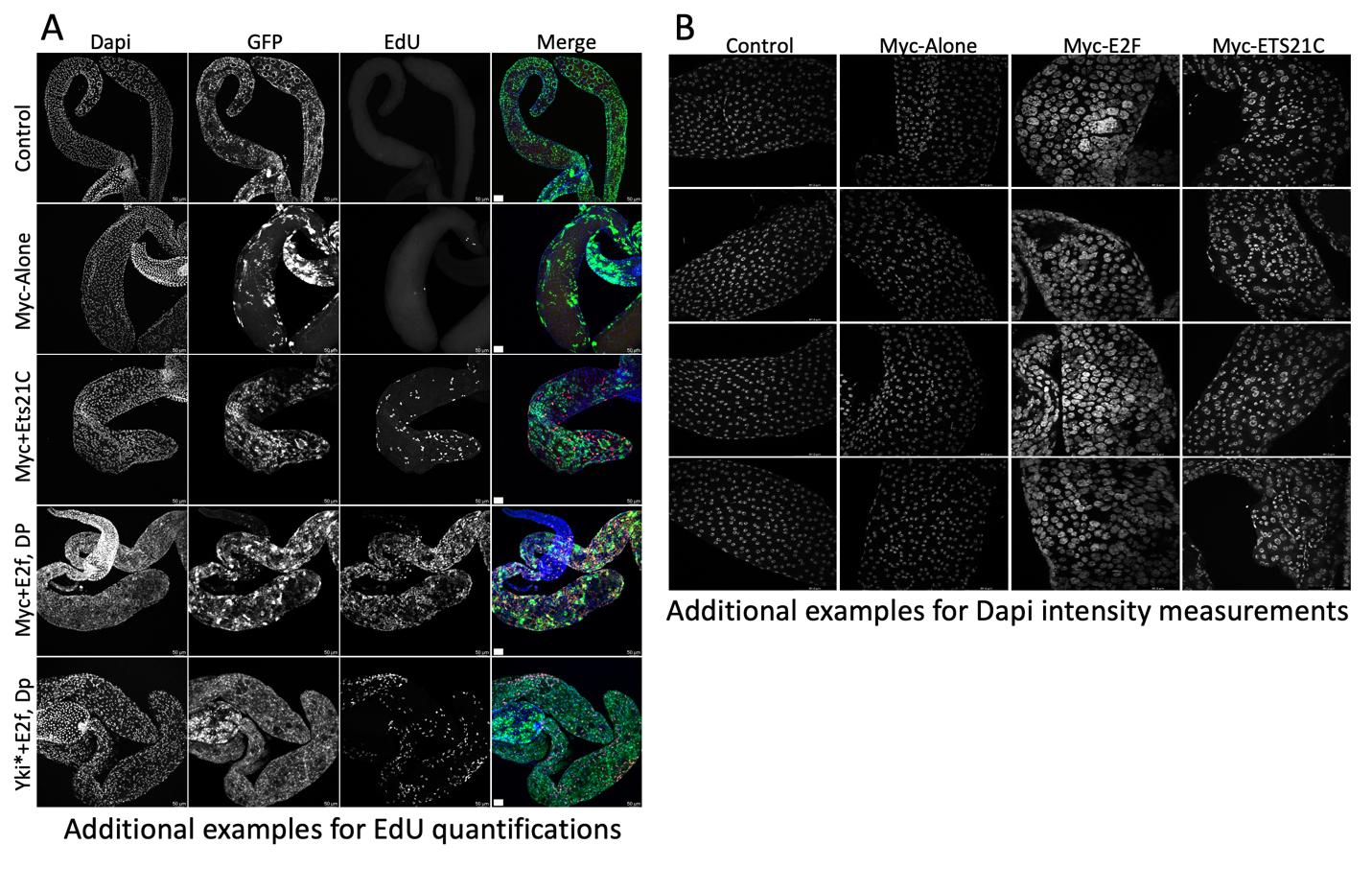

**Supp. Fig. 8 Additional examples used for EdU and Dapi intensity quantifications in Fig. 6**.

A) Samples were labeled by EdU feeding for four days from 6th-10th day of transgene expression.

B) Samples were expressing transgenes for 10-days at 25 C. These sample were used to generate the nuclei intensity graph. Fifty nuclei from each image were used to generate the graph. Fiji was used to determine the integrated Dapi intensity.

**Supplemental Methods**

### qRT-PCR primers used:

| ITGB1 | **Forward primer** | CCGCGCGGAAAAGATGAAT |
| --- | --- | --- |
|  | **Reverse primer** | CACATCGTGCAGAAGTAGGC |
| PTPN2 | **Forward primer** | CAGTGTGAAGCTCTTGTCAGA |
|  | **Reverse primer** | AGGGTTCAAGGAGCCAGATT |
| NOTCH2 | **Forward primer** | CTCAACCTGCCTGGTTCCTA |
|  | **Reverse primer** | GTGCTCCCTTCAAAACCTGG |
| CXCL8 | **Forward primer** | AAAGACATACTCCAAACCTTTCCAC |
|  | **Reverse primer** | CTCTGCACCCAGTTTTCCTTG |
| IL6 | **Forward primer** | TACATCCTCGACGGCATCTC |
|  | **Reverse primer** | CACCAGGCAAGTCTCCTCAT |

### Fold enrichment and Hypergeometric distribution (Supplement)

We compared the upregulated gene sets for fold enrichment and statistical significance via hypergeometric calculations. We used a threshold of log_2_ fold-change of 1 for greater and an adjusted p-value of 0.05 or smaller for the bulk RNA-Seq we generated. We use genes the authors identified as statistically significant and upregulated for the published work. The Python package Scipy.stats was used to calculate the hypergeometric probability number and the math package for convering the fold change to log_2_. An example code for fold enrichment and Hypergeometric Distribution calculations available in supplementary Fig. 12. In total we compared our datasets to 11 datasets from 7 published works.

| Cell polarity opposes Jak/STAT-mediated Escargot activation that drives intratumor heterogeneity in a Drosophila tumor model  Renal NF-κB activation impairs uric acid homeostasis to promote tumor-associated mortality independent of wasting  Shared enhancer gene regulatory networks between wound and oncogenic programs  Ets21c, Fos and Ftz-F1 drives JNK-mediated tumor malignancy  An Ectopic Network of Transcription Factors Regulated by Hippo Signaling Drives Growth and Invasion of a Malignant Tumor Model  Promoter Proximal Pausing Limits Tumorous Growth Induced by the Yki Transcription Factor in Drosop  The Drosophila Duox maturation factor is a key component of a positive feedback loop that sustains regeneration signaling | PMID: 36709425  PMID: 36029766  PMID: 37133250  PMID: 26398940  PMID: 27476594  PMID: 32737120  PMID: 28753614 |
| --- | --- |
